# Cell cycle alterations associate with a redistribution of mutation rates across chromosomal domains in human cancers

**DOI:** 10.1101/2022.10.24.513586

**Authors:** Marina Salvadores, Fran Supek

## Abstract

Somatic mutations in human cells have a highly heterogeneous genomic distribution, with increased burden in late-replication time (RT), heterochromatic domains of chromosomes. This regional mutation density (RMD) landscape is known to vary between cancer types, in association with tissue-specific RT or chromatin organization. Here, we hypothesized that regional mutation rates additionally vary between individual tumors in a manner independent of cell type, and that recurrent alterations in DNA replication programs and/or chromatin organization may underlie this. Here, we identified various RMD signatures that describe a global genome-wide mutation redistribution across many megabase-sized domains in >4000 tumors. We identified two novel global RMD signatures of somatic mutation landscapes that were universally observed across various cancer types. First, we identified a mutation rate redistribution preferentially affecting facultative heterochromatin, Polycomb-marked domains, and enriched in subtelomeric regions. This RMD signature strongly reflects regional plasticity in DNA replication time and in heterochromatin domains observed across tumors and cultured cells, which was linked with a stem-like phenotype and a higher expression of cell cycle genes. Consistently, occurrence of this global mutation pattern in cancers is associated with altered cell cycle control via loss of activity of the *RB1* tumor suppressor gene. Second, we identified another independant global RMD signature associated with loss-of-function of the *TP53* pathway, mainly affecting the redistribution of mutation rates away from late RT regions. The local mutation supply towards 26%-75% cancer driver genes is altered in the tumors affected by the global RMD signatures detected herein, including additionally a known pattern of a general loss of mutation rate heterogeneity due to DNA repair failures that we quantify. Our study highlights that somatic mutation rates at the domain scale are variable across tumors in a manner associated with loss of cell cycle control via *RB1* or *TP53*, which may trigger the local remodeling of chromatin state and the RT program in cancers.

## Introduction

During cancer evolution, somatic cells accumulate a number of mutations, most of them nonselected “passengers”. These somatic mutations are caused by different mutagenic processes, many of which generate higher mutation rates in late DNA replication time (RT), inactive, heterochromatic DNA. This is likely due to higher activity and/or accuracy of DNA repair in early-replicating, active chromosomal domains ^1,2^.

These chromosomal segments are defined roughly at the megabase scale, and tend to correspond to topologically associating domains (TADs) and RT domains ^3–5^. Regional mutation density (RMD) of mutations in megabase-sized domains in the human genome correlates with domain RT, local gene expression levels, chromatin accessibility (as DNAse hypersensitive sites (DHS)), density of inactive histone marks such as H3K9me3 and inversely with density of active marks such as H3K4me3 ^1,6–8^. The RMD signatures have been shown to be tissuespecific, and can be used to predict cancer type, and potentially subtype at high accuracy ^9,10^. The tissue-specificity of RMD is paralleled in the tissue-specificity of active or inactive domains. For instance, the domain that switches from late-RT to early-RT, or where genes increase in expression levels, or that gets more accessible chromatin in a particular tissue, also exhibits a reduced rate of somatic mutations in that tissue ^1,6^; this property may help identify the cell-of-origin of some cancers ^11^.

Apart from variation in active chromatin and gene expressions between tissues, recent work suggests existence of gene expression programs that are variably active between tumors originating from the same tissue (and also between individual cells), but are recurrently seen across many different tissues ^12,13^. Such programs may conceivably drive, or be driven by chromatin remodeling that activates or silences chromosomal domains. Indeed, chromatin remodeling was widely reported to occur during tumor evolution, and this can manifest as changes in RT between normal and cancerous cells, loss of DNA methylation in some chromosomal domains with cell cycling, as well as a generalized loss of heterochromatin upon transformation ^14–18^. These changes in RT, DNA methylation and heterochromatin occuring in cancer cells may plausibly affect chromosomal stability, given the links of various DNA damage and repair processes and chromatin organization ^1,2,16,19–21^.

Here, we hypothesized that chromatin remodeling that occurs variably between tumors may generate inter-individual variation in regional mutation rates, beyond the tissue identity or cell-of-origin identity effects on mutagenesis.

We study the RMD profiles at the megabase scale of somatic mutations from tumor wholegenome sequences, modeling this mutational portrait as a mixture of several underlying regional distributions, which may correspond to different mechanisms that produce or prevent mutations preferentially in some genomic domains. To disentangle these distributions, we apply an unsupervised factorization approach and extract RMD signatures from ~4200 whole genome sequenced human tumors. Some of these RMD signatures represent the expected differences between tissues/cell types, or they may represent consequences of common DNA repair failures. However others are novel and are associated with RT variation and with chromatin remodeling upon cell cycle disturbances. We characterize the differences between individuals in the usage of these different RMD distributions of mutations, suggesting that the chromatin remodeling RMD signatures are ubiquitous amongst human cancers. They associate with alterations in cell cycle genes *RB1* and *TP53* and reflect wide-spread mutation redistribution across domains and affect mutation supply to regions harboring cancer genes.

## Results

### Inter-individual variability in megabase-scale regional mutation density in human tumors

We hypothesized that, in addition to the known RMD variation between cancer types, the RMD patterns encompass variability between individual tumors that is observed independently of tissue-of-origin. To test this, we performed a global unsupervised analysis of diversity in one-megabase (1 Mb) RMD patterns across 4221 whole-genome sequenced tumors. To prevent confounding by the variable SNV mutational signatures across tumors ^22^ we controlled for trinucleotide composition across the 1 Mb windows by sampling (Methods). We additionally normalized the RMDs at chromosome arm-level to control for possible confounding of large-scale copy-number alterations (CNA) on mutation rates. Finally we removed known mutation hotspots (e.g. CTCF binding sites, see Methods), and also exons of all protein-coding genes to reduce confounding by selected mutations.

To validate the obtained tumor RMD features, we applied a Principal Component (PC) analysis on the RMD profiles across all tumor samples (n=4221). Expectedly, these PCs separated different tissues and clustering on the RMD mutational profile PCs largely reflected tissue identity (Fig S1a-e). We note cases where the similarity of RMD profiles suggested to ‘merge’ apparently similar cancer types (e.g. RMD_cluster3 with various digestive tract cancers, or the squamous-like RMD_cluster11, containing head-and-neck cancers, the non-melanoma skin cancers and some esophagus and lung cancers) (Fig S1e). Conversely, RMD profiles may subdivide some cancer types, for instance breast cancer samples in RMD_cluster6 (ovarian-like) have visually distinct RMD profiles from the typical breast-like RMD_cluster9 (Fig 1ab; Fig S1ef); the former are from the triple negative breast subtype, which has similarities to ovarian cancer by gene expression ^23^. This suggests that RMD profiles may be more generally useful for subtyping. This is illustrated in the case of head-and-neck cancer, which can be split into RMD_cluster11 (squamous-like, includes non-melanoma skin cancers) and RMD_cluster13 (this group also contains some lung cancers and so would be considered lung-like) (Fig 1ac; Fig S1e).

**Figure 1.**
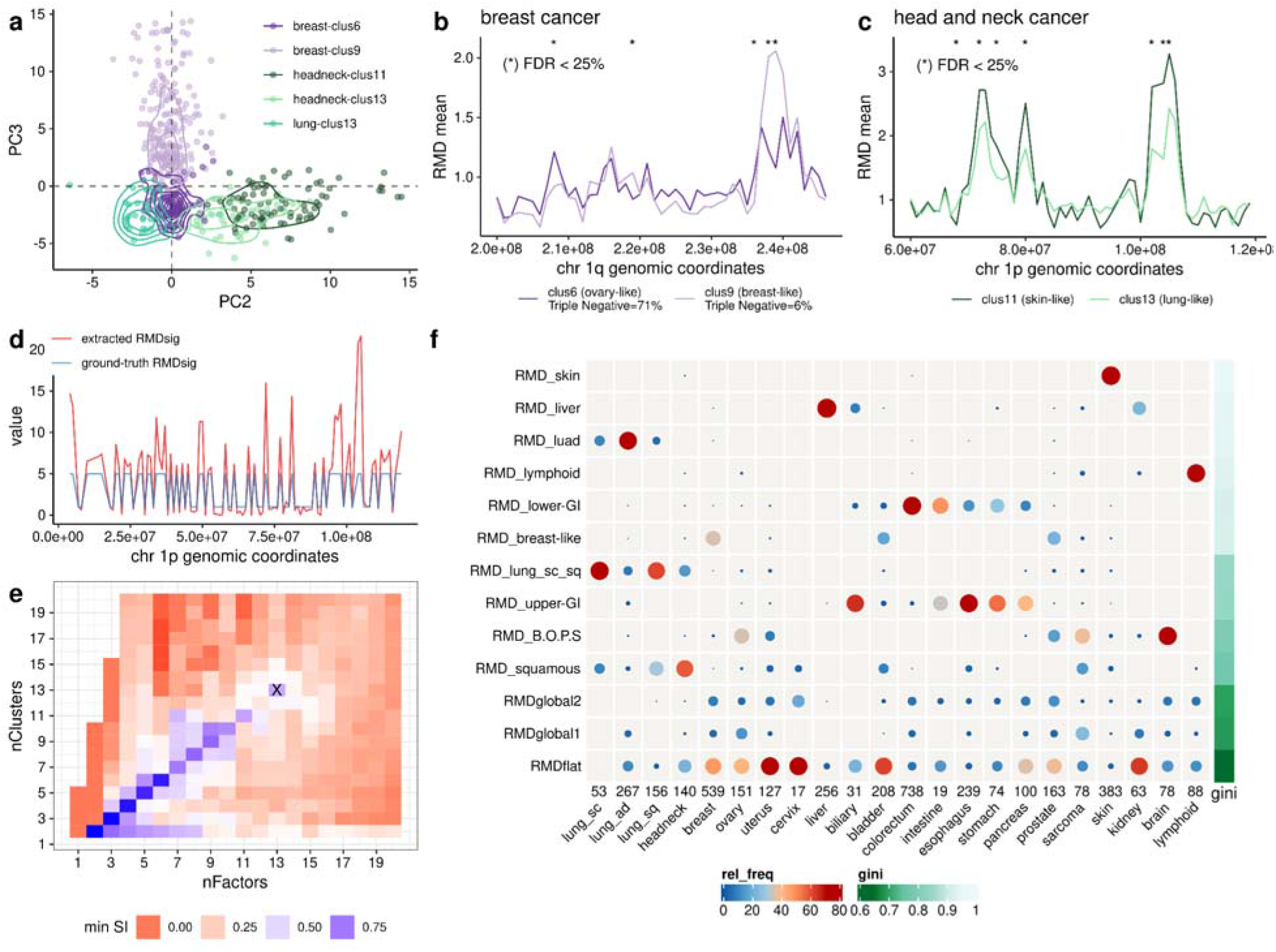
Identifying RMD signatures by an application of a NMF-based methodology to RMD of human tumors. **a)** PC2 and PC3 from a PCA in the RMD of breast, head and neck and lung human tumors separate the subgroups within the same tissue. **b)** Mean RMD profiles for breast cancer samples in cluster 6 (n = 76) and cluster 9 (n= 211), shown for chr 1q. **c)** Mean RMD profiles for head and neck squamous samples in cluster 11 (n = 81) and cluster 13 (n= 41), for chr 1p. **d)** Example signature from a simulation study, comparing window weights for an extracted NMF signature and its matching simulated ground-truth signature along chr 1p. See Supplementary Figs 2-4 for additional simulation data. **e)** NMF run on data from 4221 human tumors. Minimum silhouette index (SI) across clusters (RMD signatures) for different numbers of NMF factors and clusters. Selected case (nFactor=13, nCluster=13) is marked with a cross. **f)** Overview of the 13 RMD signatures extracted (rows) and their distribution across different cancer types (columns). The circle size and, equivalently, color corresponds to the fraction of samples from a specific cancer type exhibiting a specific signature (signature exposure >= 0.177). Total number of samples per cancer type written beneath table. The Gini index quantifies the distribution of the signature across different cancer types; higher index means more specificity to few cancer types.

As a control, our RMD features captured two known examples of regional redistribution of mutation rates: one affects specific genomic loci (somatic hypermutation regions in B-lymphocytes (Fig S1gh)), and the other causes a global homogenization (‘flattening’) of the RMDs along the genome in MMR-deficient tumors ^1^ (Fig S1g) and in APOBEC-mutagenized tumors ^24^ of various cancer types (discuss below).

Overall, even though the RMD profiles contain tissue-specific signal, there is systematic RMD variability in certain tumor genomes observed independently of the tissue-of-origin.

### A methodology to detect inter-individual variation in regional mutation density

Aiming to separate the inter-tumor RMD variability patterns presumably independent of tissue-of-origin from the tissue-specific RMD variability, we devised a methodology analogous to that recently used for extracting trinucleotide SNV mutational signatures ^22,25,26^ however here applied to 2540 megabase-sized domains instead of the typical 96-channel trinucleotide SNV spectrum. In brief, non-negative matrix factorization (NMF) is repeatedly applied to bootstrap samples of mutational counts per 1 Mb windows, normalized for trinucleotide composition as above, to find NMF solutions (sets of factors) that are consistent across bootstrap runs and thus robust to noise in the data. These solutions contain multiple RMD signatures (factors), each with RMD window weights (all 1 Mb windows with varying contributions) and RMD sample ‘exposures’ or activities (the weight of each tumor for that signature).

To test whether our NMF method is sufficiently powered to capture RMD inter-individual variability, we simulated cancer genomes containing ground-truth patterns of RMD that affected a variable number of windows, being present in variable number of tumor samples, and present at variable intensity (fold-increase over canonical mutation rate distribution at each window) (Fig S2a, see detailed description in Methods). We ran our NMF methodology for these different scenarios independently. We selected the number of factors and clusters based on the silhouette index (SI), over multiple runs of NMF (Fig S2b), and then matching the known ground-truth signatures to estimate accuracy (Methods, Fig S3). We show an example of an extracted RMD signature compared to its ground-truth signature in Fig 1d.

By comparing the different scenarios (Fig S4), encouragingly, we observed that even with a small fraction of tumor samples affected by a signature (5%), the ground-truth RMD signatures can be identified reliably, as long as the contribution of the RMD signature to the total mutation burden is reasonably high (>=20%). In addition, we observed that the NMF setup is very robust to the number of windows affected and is usually able to recover RMD signatures that affect as little as 10% of all windows. Out of other characteristics that may affect power to recover RMD signatures, we identified the signature strength/exposure (fold-enrichment) as showing the highest effect, thus the signatures with subtle effects on RMD might not be recovered (Fig S4). In summary, our simulations support that our NMF methodology can recover the genome-wide RMD signatures across chromosomal domains in a wide variety of scenarios.

### Varying degrees of tissue-specificity in RMD patterns observed across cancer types

We applied the NMF methodology to the somatic RMD profiles of 4221 tumor WGS, here requiring a minimum of 3 mutations/Mb per sample thus restricting to tumors with less noisy RMD profiles (as a limitation, we note that this may deplete some low mutation burden cancer types preferentially). In total, we extracted a total of 13 RMD signatures based on the silhouette index that scores the reproducibility of solutions upon 100 bootstraps (Fig 1ef, Fig S5).

We observed that the RMD signatures from NMF span a continuum from very tissue-specific (high Gini index, Fig 1f), to global signatures (low Gini index). We named the 10 tissue-specific RMD signatures according to the tissue or tissues they affect (e.g. RMD_upper-GI, RMD_liver), while the three global signatures that affect many cancer types were named RMDglobal1, RMDglobal2 and RMDflat (Fig 1f, Fig S5); the latter is named by the visually recognizable pattern, and also has in part known mechanisms (see below) while RMDglobal and RMDglobal2 are to our knowledge novel.

While some RMD signatures are extremely tissue-specific and capture the genomic regions with an increase of mutations only in that particular organ (e.g. skin in RMD_skin, or liver and some biliary and some kidney cancers in RMD_liver) (Fig 1f, Fig S5), many RMD signatures are observed in several cancer types which are apparently similar (Fig 1f, Fig S5). For instance, RMD_upper-GI signature is present in most esophagus, stomach, pancreas and biliary tumor samples, and some intestine tumors. The RMD_lower-GI, in turn, contains mainly the colorectal and most of the intestinal tumors, broadly consistent with the subdivision by developmental origin into the foregut (RMD_upper-GI) and the midgut/hindgut (RMD_lower-GI; Fig 1f). The RMD_squamous signature spans some squamous lung cancers, head-and-neck cancers, some bladder cancers (consistent with reports based on gene expression data ^27^), also expectedly some cervical and esophageal tumors, however surprisingly includes some sarcomas and uterus cancers suggesting a squamous-like phenotype. These commonalities in regional mutation rates probably reflect similarity of chromatin organization in the cell-of-origin of tumor types, stemming from anatomical site and/or cell type similarity, and shape the RMD profiles of those samples. Our RMD signatures support the proposed uses of RMD profiles for elucidating cell-of-origin and cancer development trajectories (e.g. metaplasia and/or invasion) ^11^ by matching to chromatin profiles.

### Three prevalent patterns of megabase-scale mutation rate variation observed across most somatic tissues

Interestingly, we identified 3 global RMD signatures, which capture the inter-individual RMD variability observed within most cancer types (Fig 1f, Fig S5).

While the profile of RMDflat captures the known flat RMD landscape (i.e. a low variation in mutation rates between domains) profile previously associated with MMR and NER pathway failures ^1,2^, and with high mutagenic activity of the APOBEC3 enzymes ^24^. Based on these known associations, which were recapitulated in our data (Fig S6) we hypothesized that additional RMDflat-high tumors (52% were explained by these known factors) may result from deficiencies in additional DNA repair pathways. Indeed we observed that deficiencies in the homologous recombination (HR) DNA repair were commonly associated with RMDflat (Fig S6b); see Supplementary Text S2 for discussion. Thus various DNA repair-related defects converge onto the RMDflat phenotype, with varying prevalence depending on the cancer type (Fig S6c).

Unlike the homogeneous pattern resulting from high RMDflat signature exposure, the RMDglobal1 and RMDglobal2 profiles have a complex pattern with their peaks appearing distributed throughout the chromosomes. We can rule out that RMDglobal1 and 2 are resulting from random noise, because (a) the silhouette index of RMDglobal1 and 2 (measuring robustness of their profile to noise that is introduced in repeated NMF runs) is comparable to the other RMD signatures, and (b) the autocorrelation of their profiles (measuring similarity in weights of consecutive 1 Mb windows) is comparable to the other, tissue-associated RMD signatures, which have a known biological basis (Fig S7ab).

As support to the pan-cancer analysis, we ran NMF for each cancer type independently, for the 12 cancer types with more than 100 genomes meeting criteria (Fig S8). All three global signatures can be found also in the per-cancer-type NMF runs (Fig S9). We found signatures in breast, lung and esophagus with a cosine similarity > 0.84 with RMDglobal1, and in colon, uterus and breast with a cosine similarity > 0.89 with RMDglobal2, supporting that the global signatures capture inter-tumoral RMD heterogeneity recurrently observed in various human somatic tissues.

### RMDglobal1 signature increases mutation rate in regions with plastic replication timing and heterochromatin

We were interested in the mechanism underlying the RMDglobal1 signature. To elucidate this, we first tried to predict RMDglobal1 signature spectrum (the one-megabase window weights) from epigenomic features previously reported to associate with megabase mutation rates (reviewed in ^28^): replication timing (RT), density of accessible chromatin (DNAse hypersensitive sites, DHS) and ChipSeq data for a variety of histone marks (Fig 2a). We first tried to predict the chromosome-wide profile of the RMDglobal1 signature using the average of each feature across many epigenomic datasets, which failed to predict (Fig 2a). Predicting RMDglobal1 from each RT/DHS/ChipSeq dataset individually and selecting the best predictor fared slightly better, with moderate associations (R^2^ ~= 0.2) for certain datasets with regional density of facultative heterochromatin (H3K27me3) and constitutive heterochromatin (H3K9me3) marks (Fig 2a), suggesting a role of heterochromatin organization in determining RMDglobal1.

**Figure 2.**
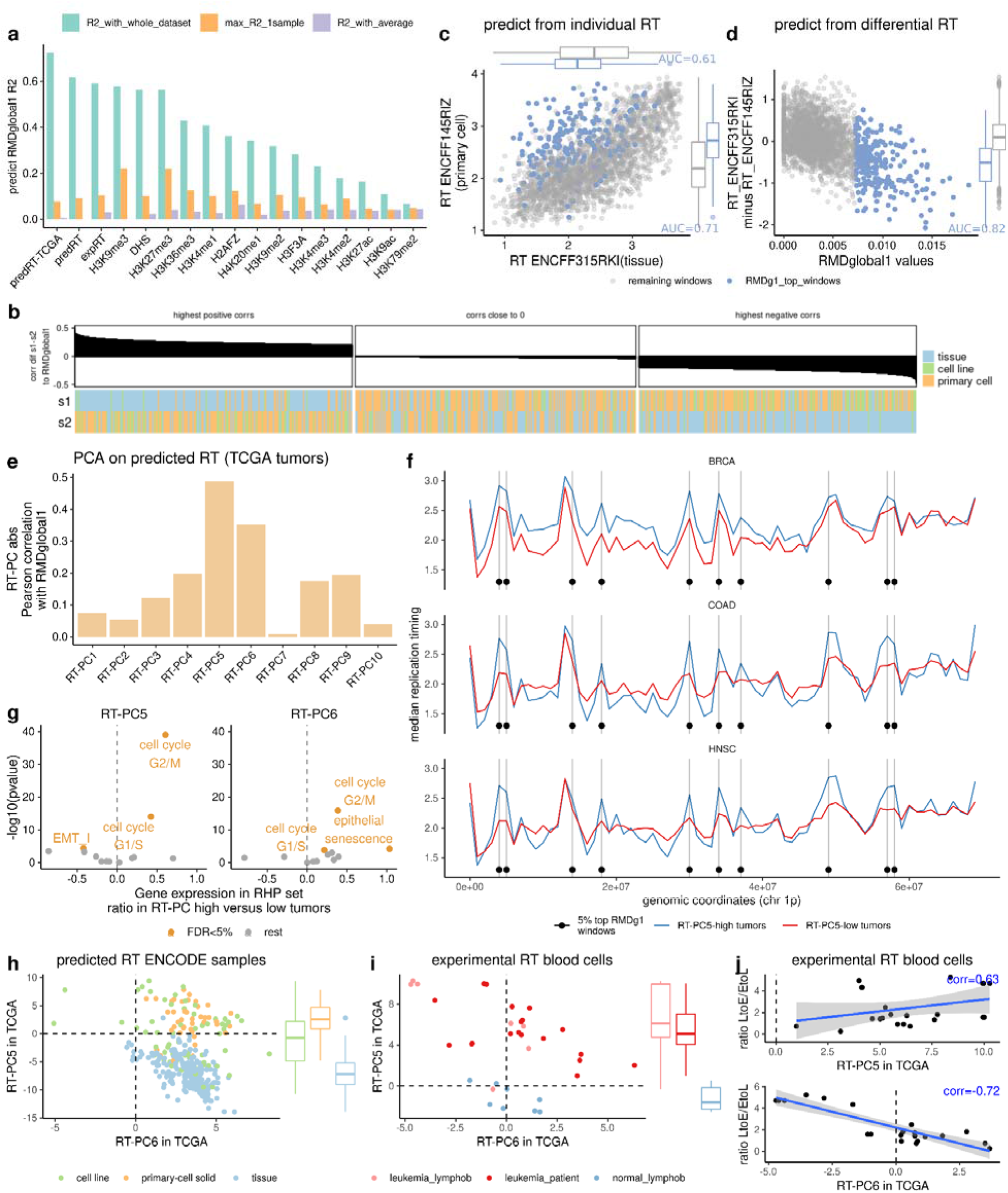
RMDglobal1 signature is linked to regional variability in replication timing. **a)** Adjusted R^2^ of a regression predicting RMDglobal1 window weights from various epigenomic features (x axis) using either the whole dataset jointly, or selecting the maximum R2 of each sample in the dataset individually, or using the average values of the feature across the samples in the dataset. **b)** Correlation between RMDglobal1 signature, and the difference between each pair of RT profiles (all combinations tested). Panel shows 1st decile (highest positive R), 5th decile (R close to 0) and 10th decile (highest negative R) deciles ordered by correlation. **c)** RT profiles for two selected samples, where dots are megabase windows, colored by their weight in RMDglobal1 signature (top decile in blue). RT of each sample individually is modestly predictive of RMDglobal1 (AUCs for discriminating top-decile windows are listed next to boxplots of RTs). **d)** Difference between the two RT profiles in panel c (on y axis) is predictive of the RMDglobal1 signature (see AUC for discriminating top-decile windows). **e)** Absolute Pearson correlation between RT-PCs and RMDglobal1 to identify the most correlated RT-PCs. **f)** Median RT profiles across samples grouped according to RT-PC5 high versus low in 3 different cancer types. Windows which are on the top 5% of RMDglobal1 weights are marked with a dot and a vertical line. **g)** Association between RT-PC5-high (top tertile) versus RT-PC5-low (bottom tertile) with the expression of genes in various RHP programs, and same for RT-PC6. **h)** Predicted RT from ENCODE data with tissues, primary cells and cell lines (predRT-ENCODE) was projected into PCs of the tumor predRT-TCGA data. **i)** Projection of experimentally determined RT data for leukemias and normal blood cells into the same PCs of predRT-TCGA data. **j)** Correlation between the projection of expRT leukemia samples in RT-PC5 and RT-PC6, and the ratio of late-to-early and early-to-late regional RT changes reported previously.

Remarkably, we observed that RMDglobal1 spectrum can be highly accurately predicted (R^2^ up to 0.7) from either RT, DHS, or the two heterochromatin marks above, however only when predicting using multiple samples jointly (contrast this with inability to predict from the averaged feature across the samples, as above (Fig 2a)). This suggests that RMDgloblal1 spectrum is explained by the variation between the samples for one chromatin feature, i.e. differences between the individual RT profiles across the genome are predictive, while the average RT profile across the genome is not. We observed the same trend using regional density of chromHMM segmentation states (Fig S10).

### An atlas of RT profiles in tumors and cultured cells links RT switching with RMDglobal1 mutation rates

Rather than histone marks, the features that best predicted RMDglobal1 were the three RT datasets (Fig 2a): (i) a collection of RT profiles from experiments [RepliChip or RepliSeq] in multiple cell types (expRT, n = 158 samples), (ii) predicted RT in a collection of noncancerous tissues, cultured primary cells and cell lines including cancer and stem cell lines (predRT, n = 597 samples), and (iii) predicted RT in human tumors (predRT-TCGA, n = 410 samples, majority measured in technical duplicate). For the latter two RT datasets, we predicted RT from DHS ^29^ or ATAC-seq data ^30^, respectively, using the Replicon tool, which infers RT profiles from local distributions in chromatin accessibility at very high accuracy ^31^ (see Methods).

Next, we aimed to characterize the mechanism of variability in RT across individuals or tissue types that predicts RMDglobal1 mutagenesis. Such RT variability appears very widespread: by calculating the difference in window-wise RT for each pair of RT samples, and correlating this difference with RMDglobal1 window weights (Fig 2b, Fig S11), we observed that the contrast of only two RT profiles can suffice to predict the RMDglobal1 spectrum using either expRT (max observed R across all pairs =0.47), predRT (max R=0.49) and predRT-TCGA (max R=0.62) datasets. We asked what types of biological samples yield such RT profiles where the contrast in RT predicts RMDglobal1 mutability. In predRT data, the best correlations are obtained when contrasting a pair that consists of one RT profile from a noncancerous intact tissue *versus* one RT profile from primary cultured cells (Fig 2b), however not when contrasting tissue to tissue, or primary cells to primary cells (Fig 2b). Conceivably, enrichment for proliferation-proficient, stemlike cells when introducing tissues into culture may alter RT, and that this altered RT is reflected in mutation rates in RMDglobal1 (see below for further discussion). As an illustrative example in a classification analysis using two selected RT profiles, one from a primary cell culture (“ENCFF145RIZ”) and one from an intact tissue (“ENCFF315RKI”), we observed that while the RT profile of each sample alone does not accurately identify RMDglobal1-high windows (Fig 2c). Remarkably, the differential RT of each window between these two RT samples can accurately classify the genomic windows with high RMDglobal1 weights (AUC = 0.82) (Fig 2d). In summary, regional switches of RT between tissues and cultured cells can predict regional somatic mutation rate switches encoded in RMDglobal1 spectrum.

### Cell cycling gene expression-associated RT in tumors is mirrored in RMDglobal1 mutagenesis

To further characterize the source of variability within RT profiles that explains RMDglobal1 signature, we applied a PCA with the predRT-TCGA dataset of RT in 410 TCGA tumors. These RT-PCs represent the archetypes of systematic variation in RT program observed across tumors, and we asked whether these local shifts in RT in tumors can explain the local changes in mutation rates we observed in tumor WGS. In particular, we correlated each RT-PC with the profile of RMDglobal1 mutation rate variation across megabase windows (Fig 2e, S12). We observed that the PCs with highest amount of variance explained either represent the average RT profile (RT-PC1, RT-PC2), or in the case of following RT-PC3 and RT-PC4 are tissue-associated RT program, separating breast from kidney and brain tumors (Fig S12ac). However, the following strongest pattern of systematic RT variation, the RT-PC5, does not exhibit a strong tissue signal, but instead correlates strongly with RMDglobal1 mutation spectrum (R=-0.49) (Fig 2e). Indeed, when we checked the RT profiles for the top RT-PC5 and bottom RT-PC5 tumors, we observed that local RT differences seen across different tissues overlap with the RMDglobal1-relevant windows i.e. those where mutation rate changes notably (Fig 2f). The next best correlation of the RMDglobal1 spectrum with RT was with RT-PC6 (R=0.35).

Thus, the RT-PC5 and 6 summarize some type of common global variation in the tumors’ RT program across cancer types, and they also predict RMDglobal1 global variation in mutation distribution. To understand the biology underlying these RT-PCs, we asked how gene expression changes between the TCGA tumors with high values of a RT-PC versus tumors with low values. We considered the “recurrent heterogeneous programs” (RHP) gene sets, representing gene expression programs that are variable in a coordinated manner between individual cancer cells, and that were recurrently observed across different cancer cell lines ^12^.

In particular, the RT-PC5 pattern correlates strongly with gene expression of RHP cell cycle genes ^12^ (there are two such sets, the G2/M and G1/S, and both correlate with RT-PC5 at p=9e-40 and 1e-14, respectively) (Fig 2g, Fig S12d), while the other RHP gene sets correlate less well with RT-PC5 (next strongest p=5e-05). We additionally also asked whether the other RT-PCs correlated with RHP cell cycle gene expression. The lower-ranking RT-PC8 and RT-PC10 did so however they (unlike RT-PC5) reflected also the epithelial-mesenchymal transition (EMT) expression program and proteasomal degradation (Fig S12d). Therefore the RT-PC5 captures specifically those global RT changes occuring in tumors in association with more rapid cell cycling but not other correlated processes. Consistently, also the RT-PC6, which did have a correlation to RMDglobal1 mutation spectrum even though more subtly, also correlates to some extent with the cell cycle RHP gene expression programs (Fig 2g, Fig S12d). Overall, this suggests that the RMDglobal1 regional mutability signature reflects the global RT program alterations associated with expression of cell cycle genes, and thus likely the variable speed of cell cycling across different tumors.

### Global RT changes observed in proliferative, cancer-like cells associate with RMDglobal1 mutation pattern

To further understand the biology of the systematic variation in RT captured by the RT-PCs relevant to mutation rates, we projected the predRT and expRT data sets originating from measurement in various tissues and cultured cells into the existing predRT-TCGA coordinate system of RT-PCs, defined by tumor RT profiles of the TCGA. Considering the predRT profiles, RT-PC5 separated tissues *versus* cultured primary cells in predRT samples (Fig 2h). One interpretation is that RT-PC5 captures the effect of tissue culture conditions on RT profiles, however we think this is unlikely because there is a considerable spread within the cultured cells group, which span across the tissue-side of RT-PC5 on the one extreme of RT-PC5 and cell line-side on the other extreme of RT-PC5 (Fig 2h). The other interpretation is that RT-PC5 captures the RT program of proliferation-capable, stem-like cells, which are normally a minority in an intact tissue, but are selected during establishment of cell culture. We favor this latter interpretation, which is consistent with the above-mentioned cell cycling RHP gene expression program association with RT-PC5 and RT-PC6. Next, considering expRT data, the RT-PC5 also separated healthy *versus* cancerous cells (here considered for blood cells, where both healthy and tumor RT measurements were available (Fig 2i)). This suggests that this property captured in RT-PC5 is more prominent in cancerous cells than in normal cells, again consistent with the property being related with cell cycling, which is often unchecked in cancer.

In summary, the RT-PC5 is a global alteration in RT program seen in tumors, which separates intact tissue samples or tumors with lower cell cycle gene expression on one side, and cultured primary cells or tumors with higher cell cycle gene expression on the other side. RT-PC5 predicts the RMDglobal1 spectrum of window weights, suggesting that the windows with higher mutation rate changes are those windows that undergo changes in RT in more proliferative samples compared to less proliferative samples.

Within a subset of the expRT data, changes in RT were studied previously^32^, reporting late-to-early (LtoE) and early-to-late (EtoL) RT changes between noncancerous samples (lymphoblastoid cell lines) and cancers (leukemias and cell lines). Their pre-calculated ratio of LtoE/EtoL strongly correlate with our RT-PC6 (R=-0.72) and RT-PC5 (R=0.63) (Fig 2j), confirming that RMDglobal1 is linked to the genome-wide changes in RT that occur in various chromosomal domains during cancerous transformation.

As further support that PCA based results are robust, we saw the same trends when we initially performed the PCA in predRT (i.e. using a mix of tissues and cell types, rather than only TCGA tumors), and then projected the expRT data into it (Fig S13). Of note, the expRT-PC that reflects developmental changes as reported earlier ^33^ does not correlate with RMDglobal1 (Fig S14a), meaning that RMDglobal1 mutagenesis pattern does not relate to embryonal-like patterns of RT.

### RMDglobal1 mutation redistribution signature associates with *RB1* loss

To identify events that may drive the changes in tumoral RT we found linked with cell cycling gene expression, we performed a genome-wide association analysis involving somatic driver events. In particular, we aimed to detect somatic copy number alteration (CNA) events and deleterious point mutations that are associated with RMDglobal1 exposure, while adjusting for cancer type and for confounding between linked neighboring CNAs (qq plots in Fig S14b; Methods for details). Here, we considered 1543 chromatin modifier genes, cell cycle genes, DNA replication and repair genes and cancer genes, compared against a background of 1000 randomly chosen control genes (Methods).

For CNA, we found a strong positive association of RMDglobal1 mutation rate redistribution with deletions of the *RB1* tumor suppressor with important roles in cell cycle control and chromatin organization (FDR=0.05%, and better p-value than all control genes) (Fig 3ab, Fig S15a). Because CNA often affects large chromosomal segments, we also checked associations with *RB1* neighboring genes (Fig 3c), noting that *RB1* is at the CNA frequency peak (by mean estimated copy-number across tumors), meaning it is the likely causal gene in the CNA segment. Strength of RMDglobal1 association with *RB1* is gene dosage dependent (Fig S15b), solidifying the causal link of *RB1* loss with mutation rate redistribution. Further, we see that the effect of *RB1* point mutations shows a trend in the same direction as the *RB1* deletions (*RB1* mutations are rarer, thus this trend is nonsignificant) (Fig S15c). As independent supporting evidence, we identified deletions in *CDK6*, a negative regulator upstream of *RB1*, as the CNA event negatively associated with RMDglobal1 with the strongest p-value, exceeding any of the control genes considered (Fig 3a).

**Figure 3.**
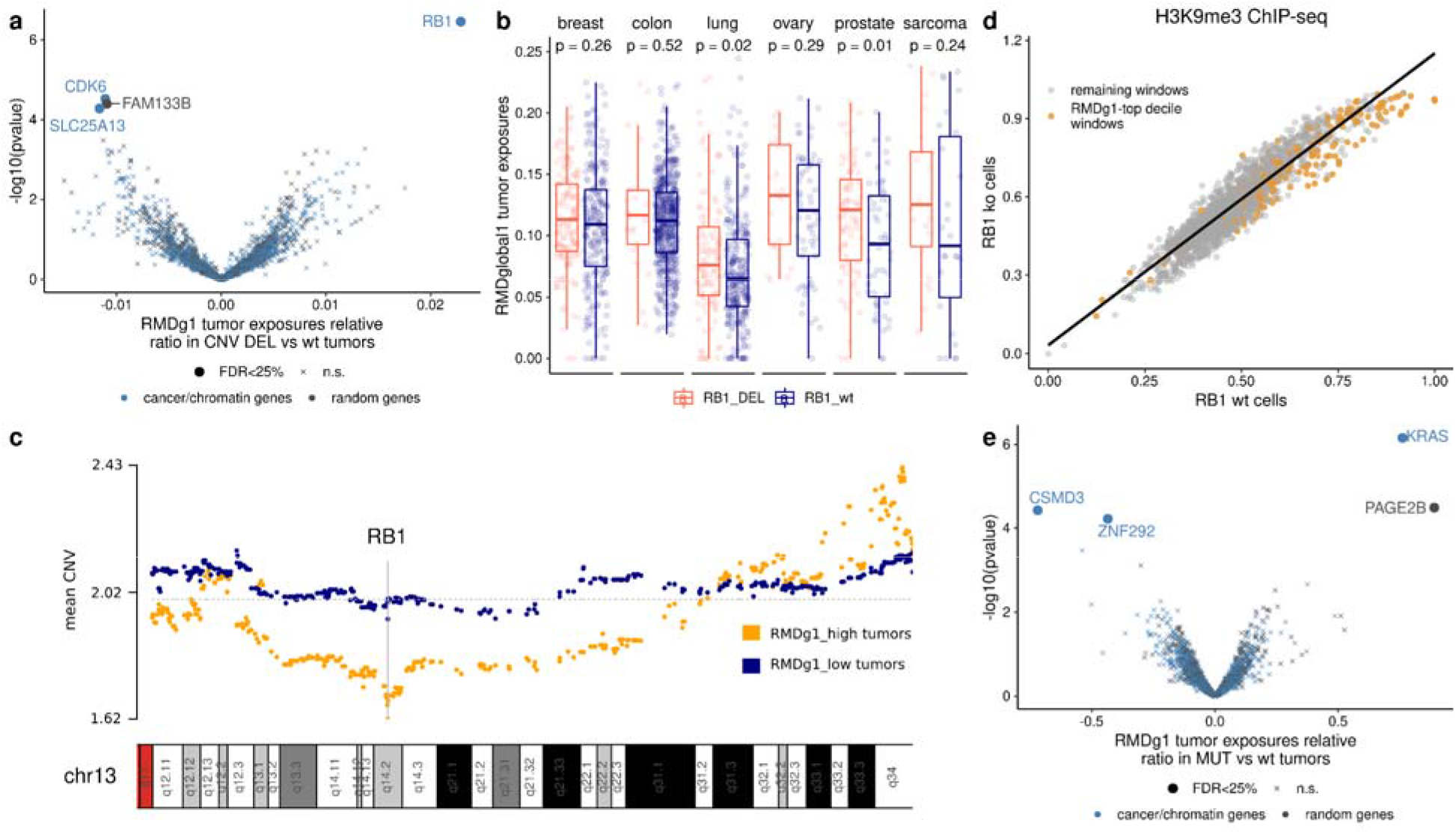
Genetic alterations associated with the activity of RMDglobal1 mutation redistribution signature. **a)** Associations between CNA deletions and tumors with higher RMDglobal1 exposures in a pan-cancer analysis, adjusting for cancer type and for global CNA patterns (Methods). N=1543 cancer genes and chromatin-related genes are shown (dots), as well as a 1000 set of randomly chosen genes (crosses). **b)** Differences in RMDglobal1 exposures between RB1 deletion (−1 or −2 deletion) and wt for several cancer types (those with the highest number of samples with *RB1* deletion); remainder in Fig S12. **c)** Mean local CN profile in groups of tumors, grouped by RMDglobal1 high and low, of the segment of chromosome 13 containing the gene *RB1*. Each dot represents one gene. **d)** Correlation between the H3K9me3 heterochromatin profiles for samples with *RB1* knock-out (“KO”) versus wild-type (“WT”). Each dot represents a window, colored by RMDglobal1 window weight top decile versus the rest of the windows. **e)** Associations between deleterious SNV and indel mutations in the same sets of genes as in panel a, and the RMDglobal1-high versus RMDglobal1-low activity of tumor samples, in a pan-cancer analysis.

In addition to its effects on cell cycle regulation, RB1 has additional important roles in chromatin organization ^19,34–36^. In specific, *RB1* deletions change heterochromatin marks H3K9me3 and H3K27me3 in regions enriched at subtelomeres, and this associates with propensity to DNA damage therein ^19^. We found these same two histone marks more highly correlated to RMDglobal1 than other tested marks (Fig 2a), and interestingly we also found that RMDglobal1 spectrum window weights are also strongly enriched approximately 5 Mb at subtelomeres (Fig 4f).

**Figure 4.**
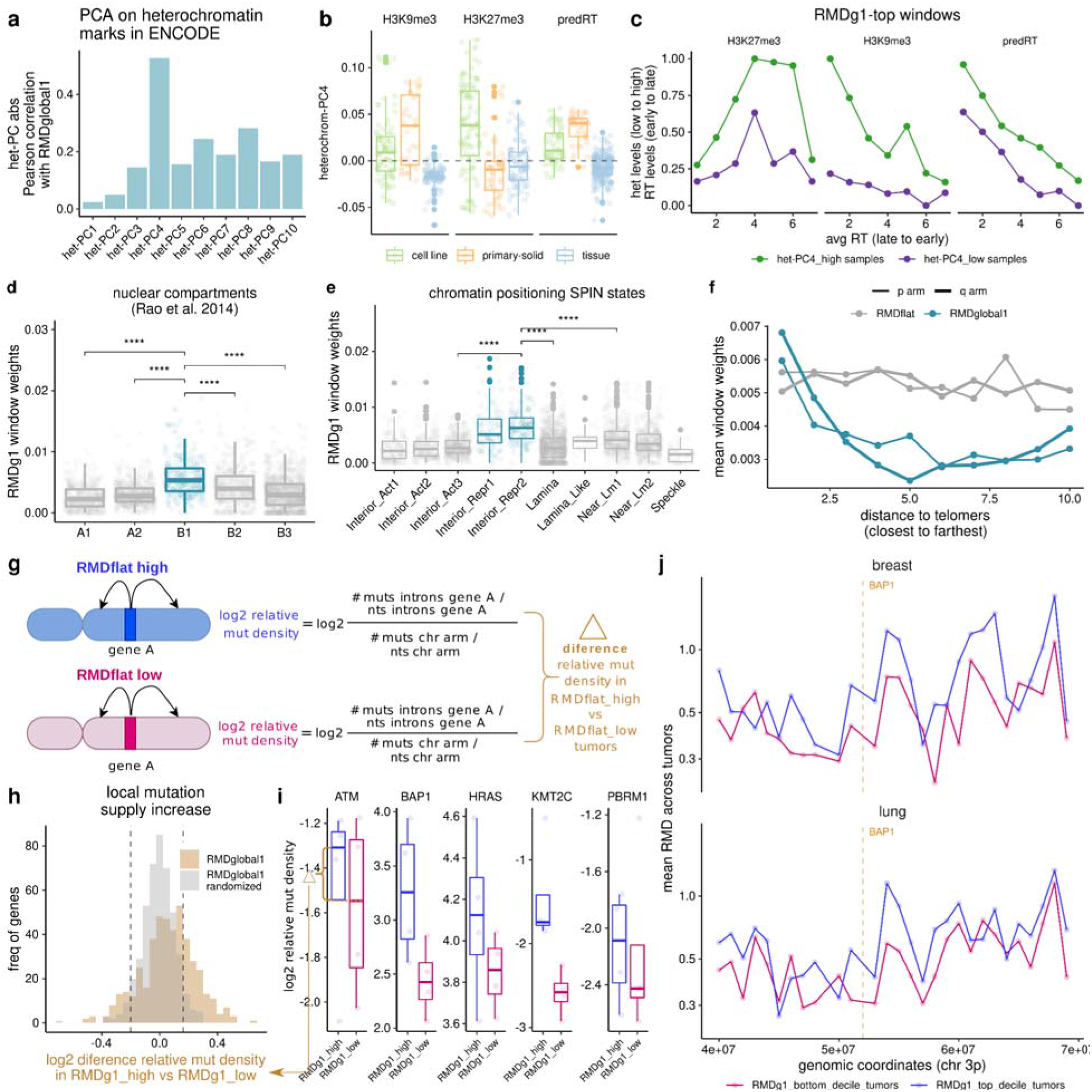
RMDglobal1 mutation rate redistribution is linked with chromatin remodeling. A PCA was performed on the predicted RT and the heterochromatin marks (H3K27me3 and H3K9me3) from ENCODE data. **a)** Absolute Pearson correlation between heterochromatin-PCs and RMDglobal1 to identify the most correlated het-PCs. **b)** Heterochromatin PC4 (selected for its high correlation with RMDglobal1 (R=0.52)) distribution across different cell types for the 3 features. **c)** Mean predicted RT, H3K27me3 and H3K9me3 across het-PC4-high versus het-PC4-low groups in ENCODE data, split by RT bins. **d)** RMDglobal1 signature across different Hi-C nuclear subcompartments from reference. **e)** RMDglobal1 signature across different SPIN nuclear compartmentalization states from ^40^. **f)** RMDglobal1 and RMDflat signature window weights compared to distance to telomeres. **g)** Schematic of the mutation supply analysis in panels h-j. **h)** Distribution for the difference in mutation density, shown for 460 cancer genes, comparing between RMDglobal1-high and low tumors, using the actual values of RMDglobal1 and as a baseline randomized of RMDglobal1. Vertical lines show 5th and 95th percentile of the randomized distribution. **i)** Mutation density for RMDglobal1-high versus low tumor samples (here, top tertile versus bottom tertile) for 5 example genes (drivers in >=4 cancer types and with the highest effect size); dots are cancer types. **j)** Mean RMD profile on chromosome 3p across the RMDglobal1-high versus low tumor groups (here, top and bottom decile by RMDglobal1), for two example cancer types. Vertical lines mark the position for the *BAP1* tumor suppressor gene (example gene in panel **i**).

### RMDglobal1 mutation redistribution affects regions that undergo heterochromatin remodelling upon RB1 loss

Prompted by the above, we asked if the location of chromatin remodelling upon *RB1* loss, considering these two heterochromatin histone marks, matches the locations of the RMDglobal1 mutation rate changes. Indeed, the changes in regional H3K9me3 profile when *RB1* is wild-type *versus* in isogenic *RB1* k.o. cells ^19^ predicted RMDglobal1 signature (adjusted R^2^=0.29), and so did changes in regional H3K27me3 albeit more subtly (adjusted R^2^=0.18) (Fig 3d, Fig S16ab). The genome regions with top 10% weights in RMDglobal1 spectrum are the regions where the level of H3K9me3 heterochromatin mark is more likely to be asymmetrically altered upon *RB1* disruption ^19^ (off-diagonal dots in Fig 3d). Overall, this overlap of heterochromatin remodelling loci upon *RB1* loss-of-function ^19^ with RMDglobal1 mutation rates change loci in tumors, further implicates RB1 activity in shaping the somatic mutation rate landscape.

We also tested associations between the presence of deleterious somatic point mutations in cancer genes and chromatin and DNA repair genes, with the ‘exposure’ to the RMDglobal1 mutagenic pattern. Here, we found the *KRAS* mutation to positively associate with RMDglobal1, at FDR=0.1% (Fig 3e, Fig S16c), and this is observed consistently across individual cancer types (Fig S16c) and significantly in colon, uterus and bladder (see Fig S16de legend for comment on lung adenocarcinoma about confounding by tobacco smoking signatures). Of note, the *KRAS* gene acts downstream of *RB1* loss-of-function with *RB1* in developmental and in tumor mouse phenotypes ^37,38^. Consistently, *KRAS* mutation and *RB1* loss (either deletion or mutation) are mutually exclusive in our tumor dataset (chi-square p < 2.2e-16), supporting that the driver alterations in *RB1* and *KRAS* may converge onto the same mutation rate redistribution phenotype, the RMDglobal1.

Motivated by these associations between RB1 loss-caused regional heterochromatin mark changes ^19^ and the RMDglobal1 regional mutation rates, we further investigated the local variation in the H3K27me3 and H3K9me3 marks across ENCODE datasets. To characterize the regional heterochromatin variation, we performed a PCA on the profiles of the two marks and the RT (predicted) together. The resulting heterochromatin-PC4 (het-PC4) correlated substantially with RMDglobal1 window weights (R=0.53) (Fig 4a). As above, the difference in the three considered features (H3K9me, H3K27me3, RT) separated the proliferative, putatively stem-like samples (het-PC4 positive) *versus* the rest of the samples (het-PC4 negative) (Fig 4b). The proliferative samples (het-PC4 positive) are later replicating in the RMDglobal1 top windows (relative increase in RT [het-PC4 high vs low] = 61%) and have higher H3K27me3 and H3K9me3 (relative increase of 55% and 78% respectively) (Fig 4c). In summary, the chromosomal domains with highest RMDglobal1 weights become later-replicating and heterochromatinized in more stem-like, proliferative cells (e.g. cell lines, primary cells), and this chromatin/RT plasticity associated with an increase of relative mutation rates in these domains.

### Gene regulation and chromatin compartments associated with the RMDglobal1 signature

The regional variation in RT and heterochromatin marks (associated with variable somatic mutation rates observed in tumors), suggests there may be concomitant changes of regional gene expression, because early RT is broadly associated with higher gene expression ^1^. Therefore we asked if there are coordinated changes in gene expression levels in certain windows between the RMDglobal1-high and RMDglobal1-low tumors. Indeed, we found several windows with coordinated gene expression upregulation and downregulation between RMDg1-high and low cancers (FDR < 25%). The coordinated downregulation windows are enriched in higher RMDglobal1 weights, compared to the windows with non-coordinated gene expression changes (Wilcoxon test, greater; downregulation p-value = 0.03; there is a also nonsignificant trend for coordinated upregulation) (Fig S17a). These regional changes in gene expression are consistent with chromatin remodeling in chromosomal domains, also mirrored in regional mutation rates (Fig S17b).

To additionally characterize the regions affected by the RT/chromatin remodelling identified herein via the RMDglobal1 mutation rates, we analyzed data from diverse genomic assays of chromatin state from various prior studies (Table S3) that reported correlations with RT. We compared the regional density of each feature with our RMDglobal1 spectrum window weights (Table S3). We noted strong correlations with Hi-C subcompartments (Fig 4d), inferred from long-range chromatin interactions at fine resolution (25 kb) ^39^. In particular, the B1 subcompartment was associated with RMDglobal1; this subcompartment replicates during middle S phase, and correlates positively with the Polycomb H3K27me3 mark (Fig S18a) but negatively with H3K36me3 suggesting that it represents facultative heterochromatin ^39^. Next, we observed a correlation with two SPIN states (Fig 4e), which were derived by integrating nuclear compartment mapping assays and chromatin interaction data ^40^. RMDglobal1 signature regions are enriched in the two “Interior repressed” SPIN states ^40^, marking regions that are inactive, however unlike other, typical heterochromatic regions these are located centrally in the nucleus, rather than peripherally (next to the lamina) ^40^. Additionally, as mentioned above RMDglobal1 important windows are enriched in subtelomeric regions (Fig 4f). In summary, the genome domains undergoing mutation rate change as per RMDglobal1 signature are enriched in the B1 facultative heterochromatin subcompartment, and in the nuclear interior located repressed chromatin.

### Mutation supply towards cancer genes is altered by global RMD signatures

Since RMDglobal1 captures a redistribution of mutation rates genome-wide, we predicted that this will affect the local supply of mutations to some cancer genes. To test this, we considered 460 cancer genes and the intronic mutation density thereof (to avoid effects of selection), further normalizing to the mutation burden of the chromosome arm to avoid effects of gross CNA on mutation rate (see Methods). We tested whether there is a difference between tumor samples with a high RMDglobal1 exposure (top tertile) versus low RMDglobal1 exposure (bottom tertile) tumors (Fig 4g-j). When compared to a randomized baseline (95th percentile of the random distribution used as cutoff); 28% of the 460 cancer genes exhibit a significant increase of mutation supply in RMDglobal1-high compared to RMDglobal1-low tumors (Fig 4h). Regarding the effect size, these genes increase mutation rates on average by 1.21-fold between the RMDglobal1-low *versus* high tertile tumors. The mutation rate density is shown for 5 example genes with a high fold-difference in Fig 4i, where for instance the median mutation rate for the *BAP1* tumor suppressor increases by 1.78-fold, for the *KMT2C* tumor suppressor by 1.79-fold, and for the *ATM* by 1.18-fold, when considering the top tertile tumors by RMDglobal1 signature of mutation redistribution.

Next, we similarly considered the RMDflat signature associated with DNA repair failures and its effects on mutability of regions containing driver genes. Tumors with RMDflat undergo an increase in mutation rates in early replicating, euchromatic regions ^24,28^. These regions also have a higher gene density, so we hypothesized that RMDflat commonly affects the mutation supply to many cancer driver genes. Remarkably, 75% of the 460 tested cancer genes ^41^ exhibit an increase in local mutation supply comparing RMDflat-low to RMDflat-high tumors, when compared to the 95th percentile of a randomized distribution (Fig S6e). The converse case was rarer: few cancer genes decreased in mutation supply in RMDflat-high tumors (9% are below the 5th percentile of the random distribution). We considered the mutation supply density for 5 example common driver genes, for which mutation supply is increased 1.8-2.5 fold between RMDflat-high and RMDflat-low tumors (Fig S6f). Considering for instance the *ARID1A* tumor suppressor gene, located in a lowly-mutated region in chromosome 1p, its mutation supply increased 1.8-fold, 2.1-fold and 2.4-fold in MSI (i.e. MMR deficient), HR deficient and APOBEC tumors (all RMDflat-high), respectively, compared to the *ARID1A* baseline mutation supply in tumors without DNA repair deficiencies (Fig S6fg). Similarly, the *BRAF* oncogene (where driver mutations are known to be highly enriched in MSI compared to MSS colorectal tumors ^42^) has considerably increased mutation supply in the RMDflat-high tumors (Fig S6fg).

### The RMDglobal2 is TP53-loss associated and reduces relative mutation rates in late replicating regions

In addition to RMDglobal1, there is a second, independently occurring mutation rate redistribution signature observed across multiple tissues, the RMDglobal2. Unlike RMDglobal1, the RMDglobal2 signature mutation rates do follow a distribution resembling the canonical RMD landscape, increasing mutation density in later RT overall, except for very late RT windows. These acquire fewer mutations than expected from their RT in the RMDglobal2 pattern (Fig 5ab). As a consequence, mutation rates increase near linearly with RT bins in tumors with high RMDglobal2, while in tumors with a low RMDglobal2 exposure the RT relationship to mutation rates is better described by a quadratic fit (Fig 5c, Fig S18b). In other words, the RMDglobal2 redistribution “linearizes” the association of RMD to RT, by suppressing the more prominent peaks in regional mutation rates, but not affecting the minor peaks and valleys.

**Figure 5.**
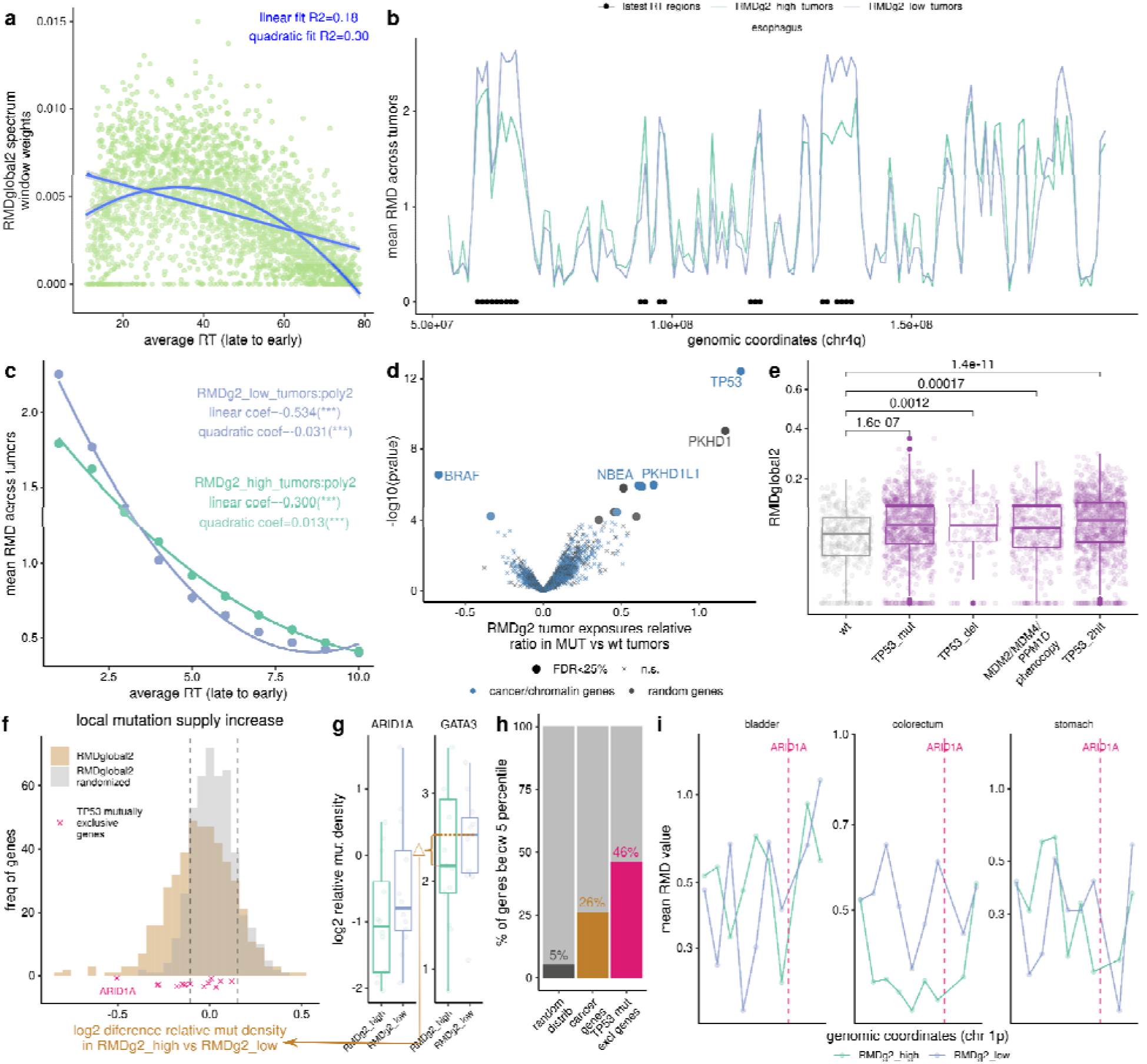
A TP53-associated mechanism underlies the RMDglobal2 mutation rate redistribution pattern. **a)** A quadratic association of RMDglobal2 signature with the average replication timing. **b)** Mean RMD profiles in chromosome 4q for the RMDglobal2-high versus low tumor samples in esophagus cancer. Latest RT windows (avRT<20) marked with black dots. **c)** Relative RMD mean profile across 10 RT bins for tumors that are RMDglobal2-high (RMDglobal2 exposures > 0.17) versus RMDglobal2-low (RMDglobal2 exposures < 0.01), showing a linearization of the link between RT and mutation rates in RMDglobal2-high. **d)** Associations between deleterious mutations in known cancer genes and chromatin-related genes (dots) and a control set of randomly chosen genes (hollow circles), and RMDglobal2 exposures in samples (p-values from Z-test on regression coefficient). **e)** RMDglobal2 signature exposures of tumor samples stratified by: wild-type for *TP53* (wt), TP53 with 1 mutation (TP53_mut), TP53 with 1 deletion (TP53_del), TP53 loss phenocopy via a amplification in *MDM2, MDM4* or *PPM1D* (TP53_pheno), or *TP53* with any two hits of the previously mentioned alteration (TP53_2hit). **f)** Distribution of the log2 difference in the relative mutation density (intronic) for 460 cancer genes, comparing between RMDglobal2 high tumors and RMDglobal2 low tumors, using the actual values (“RMDglobal2” histogram) and randomized values (“RMDglobal2 randomized” histogram). Position of the genes mutually exclusive with TP53 marked with crosses. **g)** Percentage of genes above 95 percentile of a random distribution for the random distribution, cancer genes and TP53 mutually exclusive genes. **h)** Log2 relative mutation density (normalized to flanking DNA in same chromosome arm, see Fig 4g) for RMDglobal2-high versus RMDglobal2-low for 2 example genes (TP53 mutually exclusive genes above the 95 percentile). Each dot is a cancer type. **i)** Mean RMD profile across theRMDglobal2-high versus low groups in a region of chr 1p. Vertical lines mark the position for the *ARID1A* gene.

We aimed to identify the driver event behind this redistribution of mutations away from the latest-replicating DNA domains. We proceeded to test for associations of RMDglobal2-high (top tertile) versus low (bottom tertile) tumor samples, with CNAs and deleterious mutations in cancer driver, DNA repair and chromatin modifier genes. Strikingly, we found *TP53* mutation to be uniquely strongly associated with RMDglobal2 signature (FDR = 9e-10) (Fig 5d). As supporting evidence, we found that *TP53* deletions were also positively associated (Fig 5e). Independently, the amplifications in known oncogenes that phenocopy TP53 loss (*MDM2, MDM4* and *PPM1D*) are also positively associated with the ‘exposure’ of the RMDglobal2 mutation redistribution signature (Fig 5e, Fig S19). This rules out that the *TP53* driver mutation is the consequence of the RMDglobal2 redistribution, and provides evidence for a causal effect of *TP53* pathway inactivation in the mutation rate redistribution.

Since *TP53* mutations were reported to be associated with increased burdens of CNA events ^43^, we tested whether RMDglobal2 RMD signature could be due to confounding from a multiplicity of focal CNA events, which might modify apparent local mutation rates (we note our method for RMD analysis does stringently control for confounding by arm-level CNAs, Methods). However, there is only a weak correlation between the CNA burden and RMDglobal2 signature exposure levels upon stratifying for TP53 status (R<=0.11), suggesting that RMDglobal2 largely does not reflect changes in local DNA copy number (Fig S20).

### Changes to local mutation supply because of RMD redistribution can result in epistasis-like phenomena

RMDglobal2 signature describes variation in certain genome regions, which may affect mutation supply to genes therein. We tested whether there is a difference in mutation rate in the cancer genes for RMDglobal2-high (top tertile) versus low (bottom tertile) tumor samples (Fig 5f). When compared to randomized data (5th percentile), 26% of cancer genes exhibited decreased mutation supply; only 6% genes exhibit an increased mutation supply with high RMDglobal2 (Fig 5f). As an example, we show the mutation density of *ARID1A* and *GATA3*, which decreased in mutation supply (as above, measured using intronic rates; the decrease implies they are below the 5th percentile of the randomized distribution) with high RMDglobal2 (Fig 5g). We hypothesized that apparent genetic interactions, for example mutual exclusivity with *TP53* might arise due to redistribution of mutations altering local mutation supply to genes. We thus considered 13 genes bearing mutations mutually exclusive with *TP53* mutations ^44^, hypothesizing that this might in fact be due to altered local mutation supply via RMDglobal2 in TP53 mutant cancers. Indeed we found that nearly half (6/13) of these genes were below the 5th percentile of the random distribution of local mutation rates, supporting the hypothesis (Fig 5fh). Upon inspection of the raw RMD profiles for RMDglobal2 high and low tumors for several cancer types we noted a difference in the region where *ARID1A* resides (Fig 5i). Overall, this illustrates how a global redistribution of mutation rates, here mediated by TP53 loss, can create apparent genetic interactions that may not indicate selection on functional effects of the genetic interaction. Thus, regional mutation rates, which vary extensively between tumors, should be explicitly controlled for in statistical studies of epistasis in cancer genomes.

## Discussion and concluding remarks

Mutation rates are lower in early-replicating, euchromatic DNA compared to late-replicating heterochromatic DNA ^8,45–48^. If either RT or heterochromatin (or both) are causal to mutation rates, which is likely the case and is often mediated by differential DNA repair ^1,2,6,49,50^, then local changes in RT or in heterochromatin status would change local mutation risk. Our study suggests this is commonly the case: the regions in the human genome with switching RT and switching heterochromatin status across many biological samples overlap the regions where somatic mutation rates vary across individuals and/or tumors therein (as summarized in our RMDglobal1 mutation rate redistribution signature). These regions often correspond to Polycomb-marked facultative heterochromatin residing in the B1 nuclear compartment ^39^, which is predisposed to have higher plasticity across cell types ^51^.

This locally variable RT observed across ~400 tumors is associated with cell cycle gene expression, and this locally variable heterochromatin was associated with RB1 disruption in an experimental model ^19^, plausibly reflecting various molecular consequences of accelerated and/or dysregulated cell cycles on RT and heterochromatin organization. Consistently, the changes in regional mutation rates – observed at the chromosomal domains that switch RT/heterochromatin status – are strongly associated with somatic alterations of *RB1* and *CDK6* genes in tumors. Additionally, we identify that local somatic mutation rates change appreciably due to TP53 pathway disruption in tumors, in this case most prominently in the late-replicating heterochromatic domains (as per RMDGlobal2 signature). Our data converges onto a mechanism where altered cell cycling, commonly occurring via somatic alterations of tumor suppressor genes, can trigger RT changes and heterochromatin changes, which in turn alter the local distribution of mutation rates. This affects differential mutation supply to disease genes, altering the likelihood of obtaining pathogenic mutations and steering the course of somatic evolution.

## Methods

### WGS mutation data collection and processing

We collected whole genome sequencing (WGS) somatic mutations from 6 different cohorts (Table S1). First, we downloaded 1950 WGS somatic single-nucleotide variants (SNVs) from the Pan-cancer Analysis of Whole Genomes (PCAWG) study at the International Cancer Genome Consortium ^52^ Data portal (https://dcc.icgc.org/pcawg). Second, we obtained 4823 WGS somatic SNVs from the Hartwig Medical Foundation (HMF) project ^53^ (https://www.hartwigmedicalfoundation.nl/en/). Third, we downloaded 570 WGS somatic SNVs from the Personal Oncogenomics (POG) project ^54^ from BC Cancer (https://www.bcgsc.ca/downloads/POG570/). Fourth, we obtained 724 WGS somatic SNVs from The Cancer Genome Atlas (TCGA) study as in ^9^; we used QSS_NT>=12 mutation calling threshold in this study.

Finally, we downloaded bam files for 781 WGS samples from the Clinical Proteomic Tumor Analysis Consortium (CPTAC) project ^55,56^ and bam files for 758 tumor samples from the MMRF COMMPASS project ^57^ from the GDC data portal (https://portal.gdc.cancer.gov/). Somatic variants were called using Illumina’s Strelka2 caller ^58^, using the variant calling threshold SomaticEVS >=6. Additionally, for these samples we performed a liftOver from GRCh38 to the hg19 reference genome.

We collected the samples’ metadata (MSI status, purity, ploidy, smoking history, gender) from data portals and/or from the supplementary data of the corresponding publications. Additionally, we harmonized the cancer type labels across cohorts. Here, since lung tumors in HMF data are not divided into lung squamous cell carcinoma (LUSC) and lung adenocarcinoma (LUAD) types, we used a CNA-based classifier to tentatively annotate them in the HMF data. We downloaded copy number alteration data from HMF and TCGA for lung tumor samples and adjusted for batch effects between cohorts using ComBat as described in our previous work ^59^. We trained a Ridge regression model with TCGA data to discriminate between LUSC and LUAD and applied the model to predict LUSC/LUAD in the HMF lung samples. We did not assign a label to samples with an ambiguous prediction score between 0.4 and 0.6.

Similarly, since POG breast cancer (BRCA) samples are not divided into subtypes (luminal A, luminal B, HER2+ and triple-negative) we used a gene expression classifier to annotate them. We downloaded gene expression data for TCGA and POG breast tumors and adjusted the data for batch effect using ComBat as previously described ^59^. We trained a Ridge regression model with TCGA data to discriminate between the breast cancer subtypes (one-versus-rest) and applied the model to the POG breast samples to assign them to a subtype. We did not assign 23 samples that are predicted as two subtypes and 8 that are not predicted as any subtype.

### Defining windows and filtered regions

We divided the hg19 assembly of the human genome into 1 Mb-sized windows. These divisions are performed on each chromosome arm separately. To minimize errors due to misalignment of short reads, we masked out all regions in the genome defined in the ‘CRG Alignability 75’ track ^60^ with alignability <1.0. In addition, we removed the regions that are unstable when converting between GRCh37 and GRCh38 ^61^ and the ENCODE blacklist of problematic regions of the genome ^62^.

Additionally, to minimize the effect of known sources of mutation rates variability at the subgene scale we removed CTCF binding site regions (downloaded from the Table Browser), ETS binding regions (downloaded from http://funseq2.gersteinlab.org/data/2.1.0) and APOBEC mutagenized hairpins downloaded from ^63^. Finally, we removed all coding exon regions (+-2nts, downloaded from the Table Browser) to minimize the effect of selection on mutation rates.

### Matching trinucleotide composition across megabase windows

To minimize the variability in mutational spectra confounding the analyses, we accounted for the trinucleotide composition of each window. For this, we removed trinucleotide positions from the genome in an iterative manner to reduce the difference in trinucleotide composition across windows. We selected 800,000 iterations that reach a tolerance <0.0005 (difference in relative frequency of trinucleotides between the windows). After the matching, we removed all windows that end up with less than 500,000 usable bps. The final number of analyzed windows is 2,540.

### Calculating the Regional Mutation Density (RMD) of each window

For our WGS tumor sample set (n=9,606 WGS) we counted the number of mutations in the above-defined windows. We required a minimum number of mutations per sample of 5,876, which corresponds to 3 muts/Mb (total genome = 1,958,707,652 bp). In total, 4221 tumor samples remain, which we use for the downstream analyses.

To calculate the <u>RMD</u>, we normalized the counts of each window by: (i) the nt-at-risk available for analysis in each window and (ii) the sum of mutation densities in each chromosome arm. To control for whole arm copy number alterations.

To calculate the <u>RMD applied to NMF analysis</u>, we first subsample mutations from the few hyper-mutator tumors, to prevent undue influence on overall analysis. We allow a maximum of 20 muts/Mb that is 39,174 muts. If the tumor mutation burden is higher we subsample the mutations to reduce it to that maximum value. Then, as above, we normalized the RMD by: (i) the nt-at-risk in each window [RMD = counts * average_nt_risk / nt_at_risk] and (ii) the sum of mutation density in each chromosome arm [RMD * row_mean_WG / rowMeans by chr arm]. We multiply by the average nucleotides at risk and the mean whole genome to maintain the values range of each sample for the bootstrapping.

### Applying NMF to extract RMD signatures

We applied bootstrap resampling (R function UPmultinomial from package sampling) to the RMD scores that we calculated for NMF as above. The result for each tumor sample is a vector of counts with a tumor mutation burden close to the original one but normalized by the nucleotides at risk by window and for the possible chromosome arm copy number alterations (CNA). Then, we applied NMF (R function: nmf) to the bootstrapped RMD matrices, testing different values of the rank parameter (1 to 20), herein referred to as nFact.

We repeated the bootstrapping and NMF 100 times for each nFact. We pooled all the results by nFact and performed a k-medoids clustering (R function pam), with different number-of-clusters k values (1 to 20). We calculated the silhouette index value, a clustering quality score (which here measures, effectively, how reproducible are the NMF solutions across runs), for each clustering to select the best nFact and k values.

Additionally, we also applied the same NMF methodology to each cancer type separately (n = 12 cancer types that had >100 samples available).

### Simulated data with ground-truth RMD signatures

For each cancer type, we calculated a vector of RMD values (i.e. regional mutation density mean of all samples from that cancer type) based on observed data, and super-imposed simulated ground-truth signatures onto these cancer type-derived canonical RMD patterns. We generated 9 simulated ground-truth RMD signatures with different characteristics, varying the number of windows affected by the signature (10, 20 or 50% of 2540 windows total) and the fold-enrichment of mutations in those windows (x2, x3 or x5) over the RMD window value in the canonical RMD pattern for that tissue.

In particular, we tested 9 different scenarios, varying the signature contribution to the total mutation burden (10, 20 or 40%) and the number of tumor samples affected by the signature (5, 10 or 20%). We randomly assigned the ground-truth signatures to be super-imposed onto each tumor sample (e.g. sample A will be affected by RMD signature 1 and 3 while sample B will be affected by signature 4). In total, we have simulated genomes for 9 different scenarios (different RMD signature contributions and number of tumor samples affected), each of them containing the 9 simulated ground-truth RMD signatures.

We applied the NMF methodology for the 9 different scenarios independently and obtained NMF signatures. For each case, we selected an NMF nFact and k-medoids clustering k, based on the minimum cluster silhouette index (SI) quality score. To assess the method, we compared the extracted NMF signatures with the ground-truth simulated signatures. In particular, we considered that an extracted NMF signature matches the ground-truth simulated signatures when the cosine similarity is >=0.75 only for that ground-truth simulated signature and < 0.75 for the rest.

### Analysis of differential mutation supply towards cancer genes

For 460 cancer genes from the MutPanning list ^41^ (http://www.cancer-genes.org/), we tested if they are enriched in intronic mutations in tumor samples with high RMDflat, RMDglobal1 or RMDglobal2. An enrichment will mean that there is a higher supply of mutations in the intron regions of those genes when the RMDsignature is high. For this, we considered the counts of mutations in the intronic regions of the gene, normalized to the number of mutations in the whole chromosome arm, comparing pooled tumor samples with RMD signatures high or low, by tissue. Note that the possibly different number of eligible nucleotides-at-risk in the central window, nor the length of the flanking chromosome arm are relevant in this analysis, because they cancel out when comparing one group of tumor samples (split by the RMD signature) to another group of tumor samples. We binarized the tumor samples by RMDflat, RMDglobal1 and RMDglobal2 by dividing each of them into tertiles, and keeping 1st tertile *versus* 3rd tertile for further analysis. We applied a Poisson regression with the following formula:

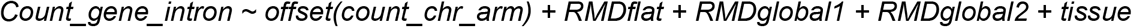

where “count” refers to mutation counts. By including the tissue as a variable in the regression, we controlled for possible confounding by cancer type. The log fold-difference in mutation supply between RMD signature high versus low tumor samples is estimated by the regression coefficients for RMDflat, RMDglobal1 and RMDglobal2 variables. As a control, we repeated the exact same analysis but randomizing the tertile assignment for the three RMD signatures prior to the regression.

### Association analysis of gene mutations with RMD global signatures

We created a subset of 1543 relevant genes: cancer genes from the MutPanning list ^41^ and Cancer Gene Census list ^64^, and furthermore we included genes associated with chromatin and DNA damage ^65^. As control, we used a subset of 1000 random genes selected as in ^65^.

We applied the analysis for two different features: copy number alterations (CNA) and deleterious point mutations. For CNA, we use the CN values by gene, using a score of −2, −1, 0, 1 or 2 for each gene. We considered a gene to be amplified if CNA value was +1 or +2 and deleted if the CNA value was −1 or −2. For deleterious mutations, we selected mutations predicted as moderate or high impact in the Hartwig (HMF) variant calls, (https://github.com/hartwigmedical/hmftools). We binarized the feature into 1 if the sample has the feature (CNA, or deleterious mutations present) or 0 if it has not. We considered CNA deletions and amplifications as two independent features. We binarized RMDflat, RMDglobal1 and RMDglobal2 by dividing each of them in tertiles and comparing tumor samples in 1st tertile *versus* 3rd tertile, by tissue.

We fit a linear model to test whether the binary genetic feature (amplification CNA, deletion CNA or deleterious mutation in a particular gene) can be explained by the RMD signatures activity being high *versus* low (i.e. upper tertile versus lower tertile). We controlled for tissue by including it as covariate. The regression formula was:

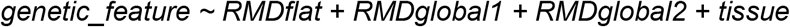

We used the regression coefficients, and p-values (according to the R function “summary”) from the variables RMDflat, RMDglobal1 and RMDglobal2 to identify genetic events associated with high levels of each RMD global signatures, suggesting possible RMD signature generating events. In the case of CNAs, to adjust for the linkage between CNA resulting in confounding, we added to the regression the PCs from a PCA on the CNA landscape across all genes. We calculated the lambda (inflation factor) for the p-value distribution of associations, while including PCs from 1 to 100 to decide the best number of PCs to include so as to minimize lambda. We included the first 55 PCs for the deletion CNA and the first 63 PCs for the amplification CNA association study.

### Epigenomic and related data sources

#### ENCODE data

We downloaded from ENCODE (https://www.encodeproject.org/) all data available for *Homo sapiens* in the genome assembly hg19 for DHS, H3F3A, H3K27me3, H3K4me1, H3K4me3, H3K9ac, H3K9me3, HiC, DNA methylation (WGBS), H2AFZ, H3K27ac, H3K36me3, H3K4me2, H3K79me2, H3K9me2 and H4K20me1 marks. Data is described in Table S2. For each of these features, we downloaded the narrow peaks, calculated their weighted density for each 1Mb window as the width of the peak multiplied by the peak value.

#### ChromHMM chromatin states

We downloaded the 25 ChromHMM states segmented files (“imputed12marks_segments”) for the 129 cell types available from Roadmap epigenomics ^66^(http://compbio.mit.edu/ChromHMM/). We calculated the density of each state for each 1Mb window as the fraction of the window covered by the chromatin state.

#### Other epigenomic data

We downloaded RT variability genomic data describing RT heterogeneity ^67^, Constitutive and Developmental RT domains ^68^, RT changes upon overexpression of the oncogene *KDM4A* ^69^, RT signatures of replication stress ^70^, RT signatures of tissues ^33^, RT states ^71^, changes in RT upon *RIF1* knock-out ^72^ and RT changes due to RT QTLs ^73^. In addition, we downloaded data for variability in DNA methylation ^15,74^, HMD and PMD regions ^16^, CpG density, gene density, lamina associated domains (LADs), asynchronous replication domains ^75^, early replicating fragile sites ^76^, SPIN states ^40^, A/B subcompartments ^39^, DHS signatures ^77^ and H3K27me3 and H3K9me profiles for *RB1* wild-type and knock-out ^19^. Data described in Table S3. We calculated the density for each feature for each 1 Mb window, and correlated this with the RMDglobal1 signature windows weights.

### Replication timing data sources and generation

We downloaded experimental RT data, from RepliChip or RepliSeq assays, from the Replication Domain database (https://www2.replicationdomain.com/index.php) ^68^ in multiple human cell types (n = 158 samples). In addition, we predicted RT using the Replicon software ^31^ from two type datasets: (i) in noncancerous tissues, cultured primary cells and cell lines including cancer and stem cells (n = 597 samples) using the DHS chromatin accessibility data downloaded from ENCODE; and (ii) in human tumors (n = 410 samples, most of them with technical replicates) using ATAC-seq data of TCGA tumors downloaded from ^30^. We used Replicon tool with the default settings.

### Analysis of coordinated gene expression changes

For the genomes from the HMF data set, we downloaded gene expression data (as adjusted TPM values) from Hartwig ^53^, available for a subset of samples for which we derived the RMD signatures. In total, we had gene expression data for 1534 samples and 18889 protein coding genes. We tested whether the gene expression values of the genes within one window show an increase or decrease compared to their flanking windows in RMDglobal1 high (exposure >= 0.13) versus RMDglobal1 low (exposure < 0.06) tumor samples using a linear regression model:

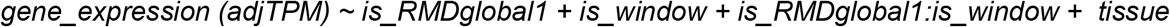

In this analysis, we removed from the datasets samples with high RMDflat or with high RMDglobal2 value (exposure > 0.15). We used samples from breast, colorectum, lung, ovary and skin because they had >=5 samples in both categories (RMDglobal1 high and low). To analyze the coordinated changes in gene expression we checked the coefficient and p-values of the interaction term *is_RMDglobal1:is_window*.

For the genomes from the TCGA data set, we downloaded gene expression data (as TPM values) from the Genomic Data Commons data portal (https://dcc.icgc.org/pcawg) for the same TCGA samples for which we predicted RT. In total, we have gene expression data for 399 overlapping samples and 20092 genes. We compared the gene expression between RT-PC5 (and RT-PC6) high and low for a group of pathways which has been reported to be related with recurrent heterogeneity across cell types ^12^ using a regression model. We binarized RT-PC5 (and RT-PC6) by dividing each into tertiles and keeping the samples in the 1st tertile to be compared versus the samples in the 3rd tertile. We applied a regression for all the genes in each RHP gene set separately. We controlled for tissue by including it as covariate. The regression formula is:

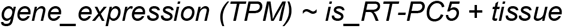

We considered the regression coefficient and its p-value of the variable is_RT-PC5. We applied the same analysis for RT-PC6.

### Clustering of RMD profiles

For RMD profiles we applied a PCA to the centered data, where rows were tumor samples and the columns were megabase windows. Next, we applied a clustering on the PC1 to PC21 using the R function tclust for robust clustering. We tested different numbers of clusters and alpha value (number of outliers removed). In addition, we tested the clustering using all PCs (PC1 to PC21) and without PC1 (PC2 to PC21), selecting the clustering for k=18 and alpha = 0.02 without PC1 based on the log likelihood measurement.

## Acknowledgements

M.S. was funded by a FPU fellowship of the Spanish government, Ministry of Universities. Work in the lab of F.S. is supported by an ERC StG “HYPER-INSIGHT” (757700), Horizon2020 project “DECIDER” (965193), Spanish government project “REPAIRSCAPE”, CaixaResearch project “POTENT-IMMUNO” (HR22-00402), an ICREA professorship to F.S., the SGR funding of the Catalan government, and the Severo Ochoa centers of excellence award of the Spanish government to the hosting institution.

This publication and the underlying research are partly facilitated by Hartwig Medical Foundation and the Center for Personalized Cancer Treatment (CPCT) which have generated, analysed and made available data for this research. In addition, data used in this publication were generated by the Clinical Proteomic Tumor Analysis Consortium (NCI/NIH). We acknowledge that the results published here are in part based upon data generated by the TCGA Research Network: http://cancergenome.nih.gov/.

