## Supplementary Figures S1-S20 for "Cell cycle alterations associate with a redistribution of mutation rates across chromosomal domains in human cancers"

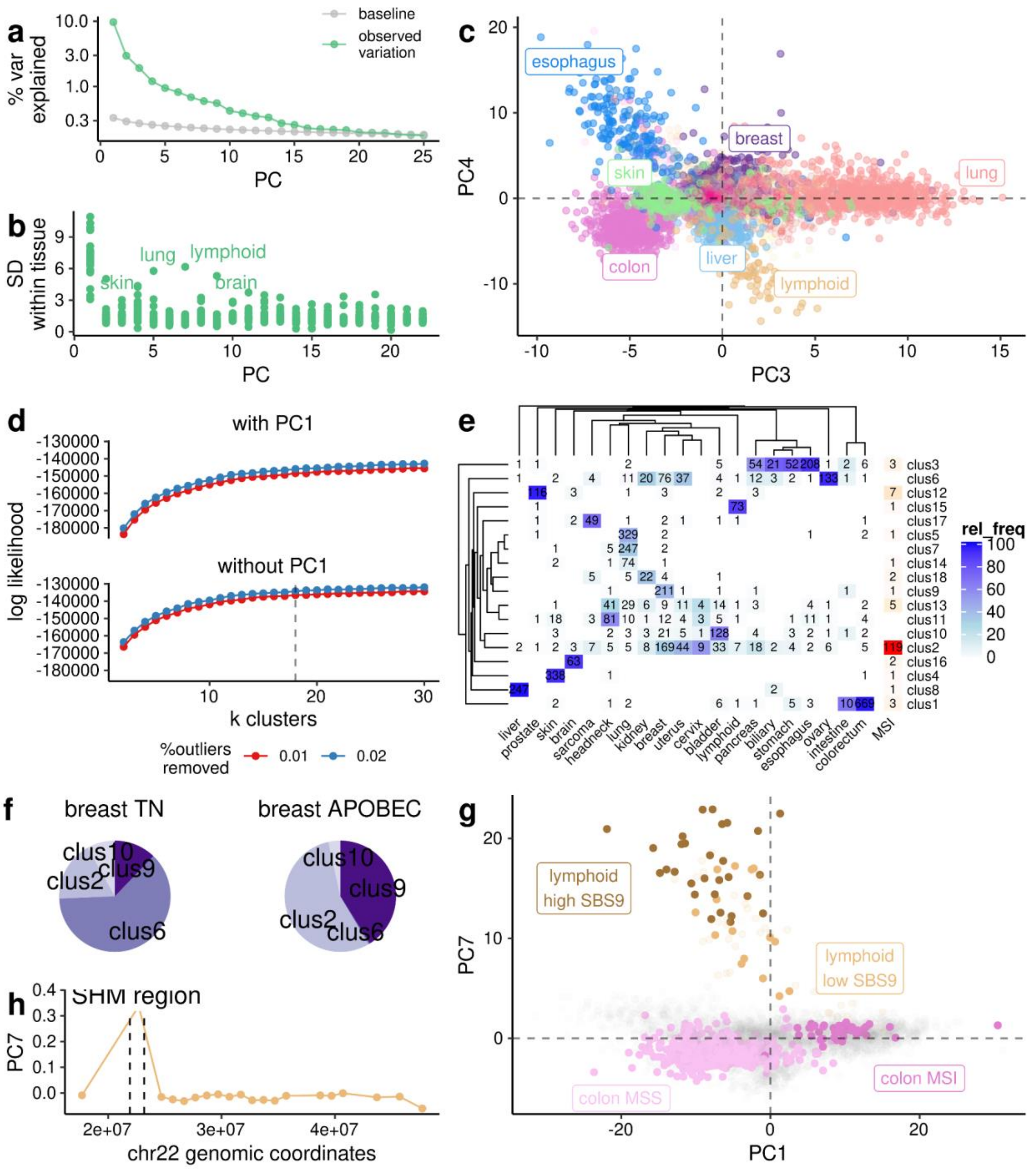


**Fig S1. RMD variability across tissues and individuals. a)** Variance explained for the first 25 PCs of a PCA on the RMD matrix (4221 samples x 2542 one-megabase windows), and a baseline (by the broken stick rule). **b)** Inter- individual variability (standard deviation) within each tissue for the first 22 PCs. **c)** PC3 and 4 separate various cancer types. **d)** Log likelihood of the clustering of the first 22 PCs using different numbers of k clusters for two cases (including and excluding PC1). **e)** Number of tumor samples from each cancer type that are assigned to each RMD-based cluster (Methods). **f)** Cluster assignment for breast cancer samples of triple negative (TN) subtype and samples with high APOBEC (>25% of mutations are in APOBEC contexts). **g)** As controls, the PC1 separates MSI versus MSS tumors, and PC7 separates lymphoid samples according to their level of the SHM mutational signature (SBS9). **h)** PC7 window weights for chromosome 22 agree with the known SHM region.

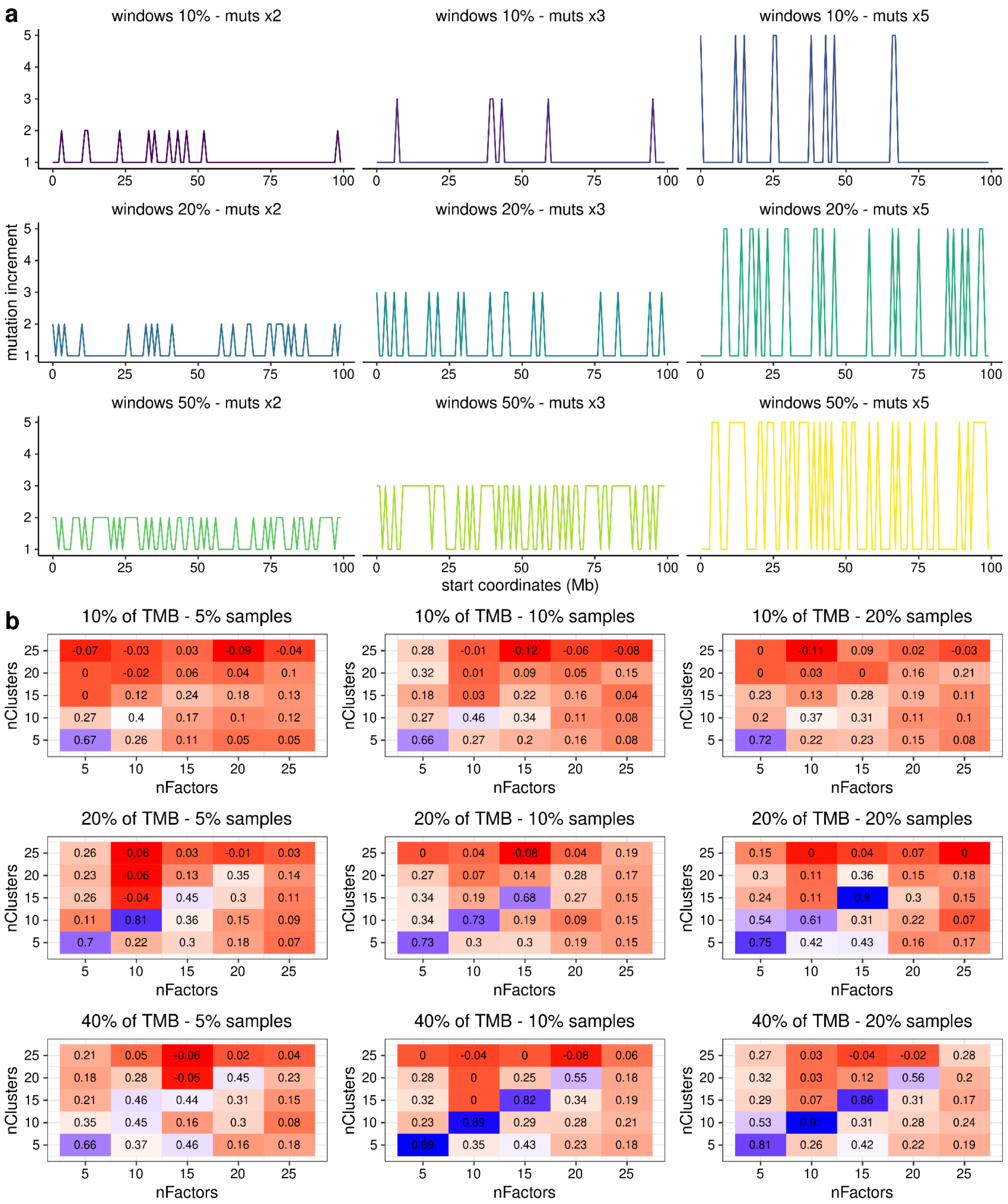


**Fig S2. Simulated ground-truth signatures to benchmark the NMF methodology. a)** Profiles for the 9 simulated signatures generated (first 100 windows of chr1). The value is the increase of mutations it will generate when multiplied by the cancer type vector (x2,x3 or x5). The windows affected by the increase could be 10, 20 or 50%. Each combination is explained in the title: sig_[windows affected]_[increase]. **b)** Min silhouette index (SI) for the clustering of the extracted signatures comparing different ks and nFacts. Each heatmap shows the min SI for a different condition varying the signature contribution to the TMB and the number of samples affected (specified in the title).

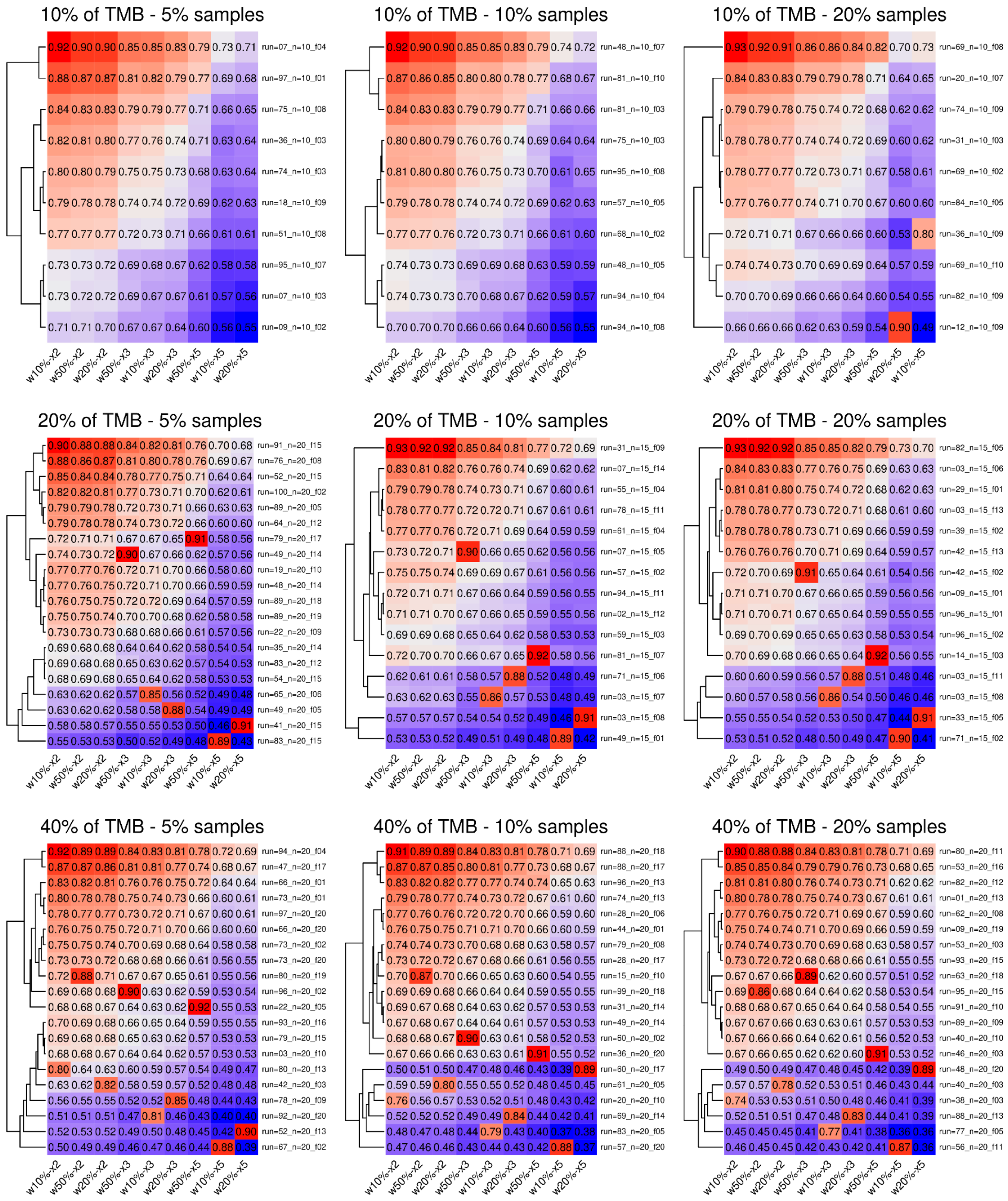


**Fig S3. Identify NMF signatures matching the ground-truth simulated signatures.** Cosine similarity values between the simulated signatures (columns) and the extracted signatures from the NMF (rows). We consider a signature to be recovered when it matches only one simulated signature with cosSim >= 0.75.

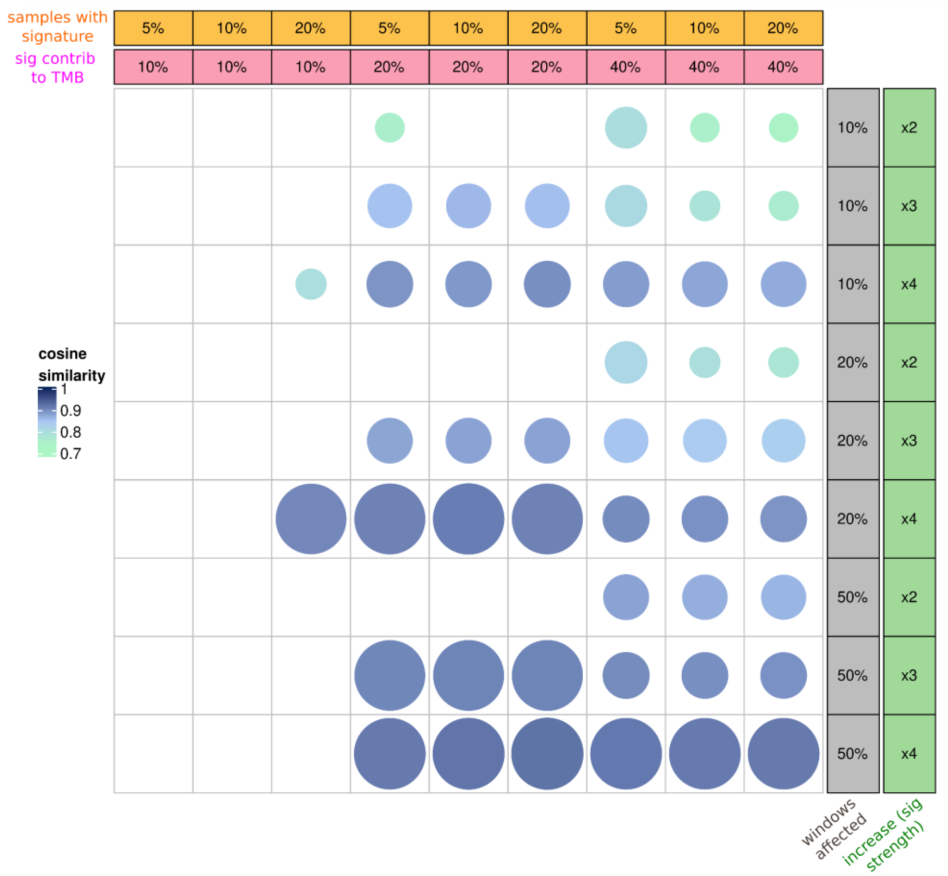


**Fig S4. Summary of the benchmarking of the NMF methodology using simulated cancer genes.** Comparison of the number of signatures recovered by the NMF method depending on the 9 conditions (columns) and the characteristics of the 9 simulated signatures (rows). We can see which simulated signatures are more often recovered (with a circle) and how strong (color and size of the circle) and under which conditions.

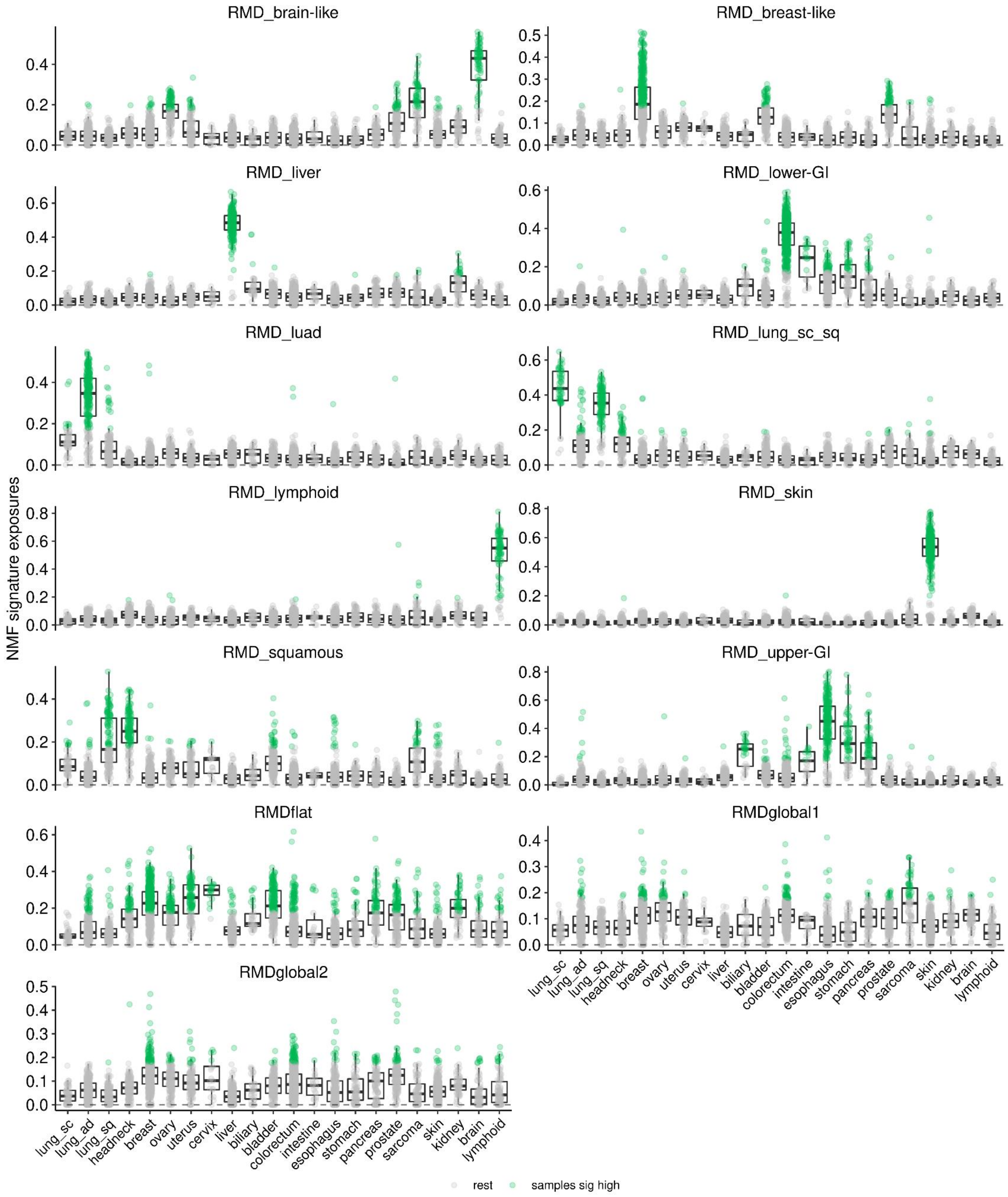


**Fig S5. The 'exposures' (activity levels) of the NMF-derived RMD signatures across various human tumor types.** Sample exposures distributed across cancer types for the 13 signatures extracted. Marked in green samples with high contribution from the signature. The threshold for considering a signature high is the sample exposure value for which 95% of the MSI samples are recovered in the RMDflat signature.

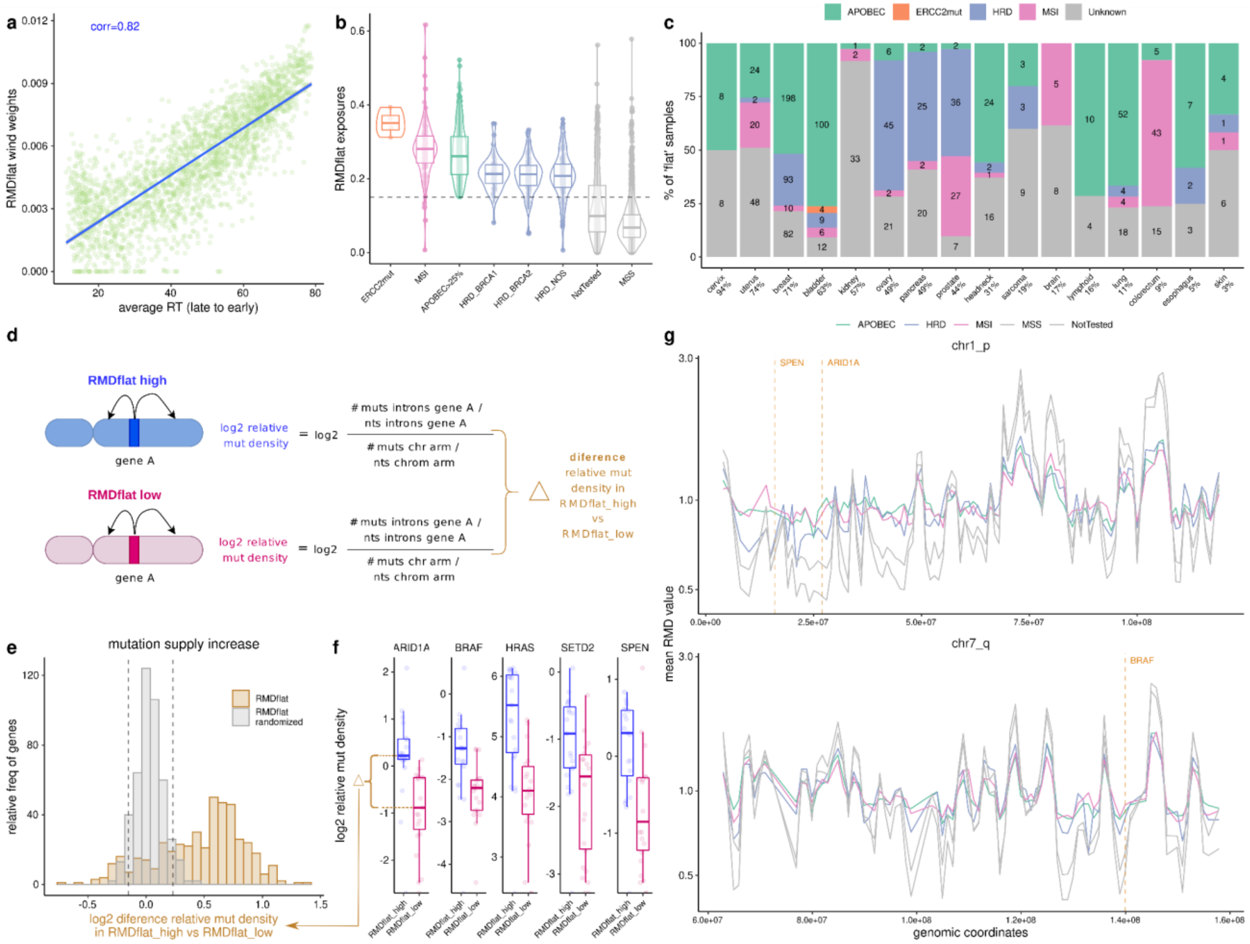


**Fig S6. Characterization of the RMDflat RMD signature, which represents a loss of mutation rate heterogeneity.** **a)** Correlation between RMDflat signature NMF window weights and the DNA replication timing (RT) (Repli-Seq average across 10 cell lines). **b)** RMDflat signature exposures (i.e. activities) for groups of tumor samples with various DNA repair failures (MSI, microsatellite instable, indiciating MMR failure; HRD, deficient homologous recombination, by the BRCA1 type or BRCA2 type or not otherwise specified), or high levels of APOBEC mutation signatures. **c)** Percentage of tumor samples with 'flat' mutation rate landscapes (RMDflat exposure>0.177, a threshold that recovers 95% of MSI samples) belonging to each of the DNA repair categories, stratified by cancer type. The percentage of 'flat' samples in each cancer type is indicated in x-axis labels. **d)** Schematic of the mutation supply analysis in panels e-g. **e)** Distribution of the log2 difference in the relative mutation density (intronic) for 460 cancer genes, comparing between RMDflat high tumors and RMDflat low tumors, using the actual values ("RMDflat" histogram) and randomized values ("RMDflat randomized" histogram). **f)** Log2 relative mutation density (normalized to flanking DNA in same chromosome arm, see panel **d**) for RMDflat-high versus RMDflat-low for 5 example genes (common drivers across >=4 cancer types and with highest effect sizes in this test) . Each dot is a cancer type. **g)** Mean RMD profile across the DNA repair groups, shown examples for chr 1p and chr 7q. Vertical lines mark the position for three example genes from panel **f**.

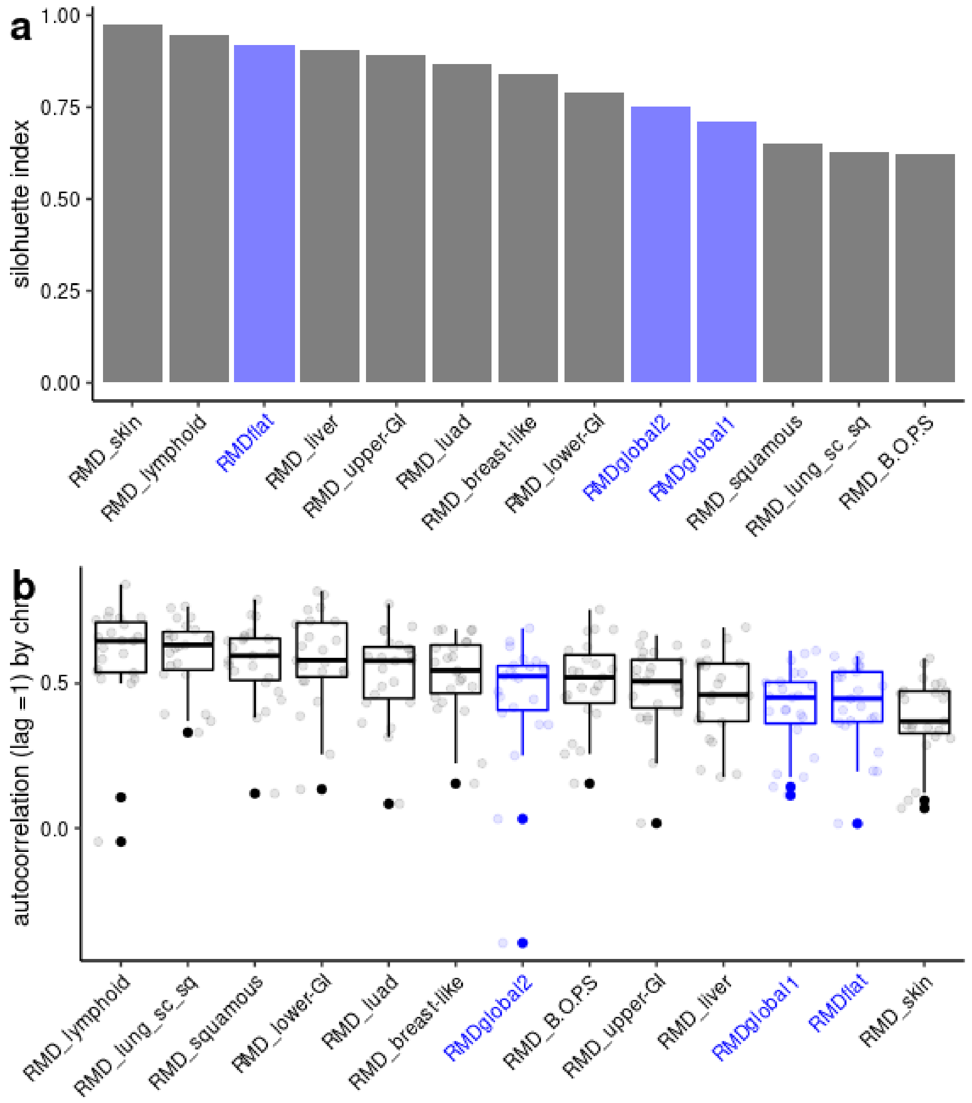


**Fig S7**. **Quality control for the extracted RMDsignatures. a)** Silhouette index value for the clustering of each RMD signature. **b)** Autocorrelation values with lag = 1 (calculated for each chromosome separately) for each RMD signature.

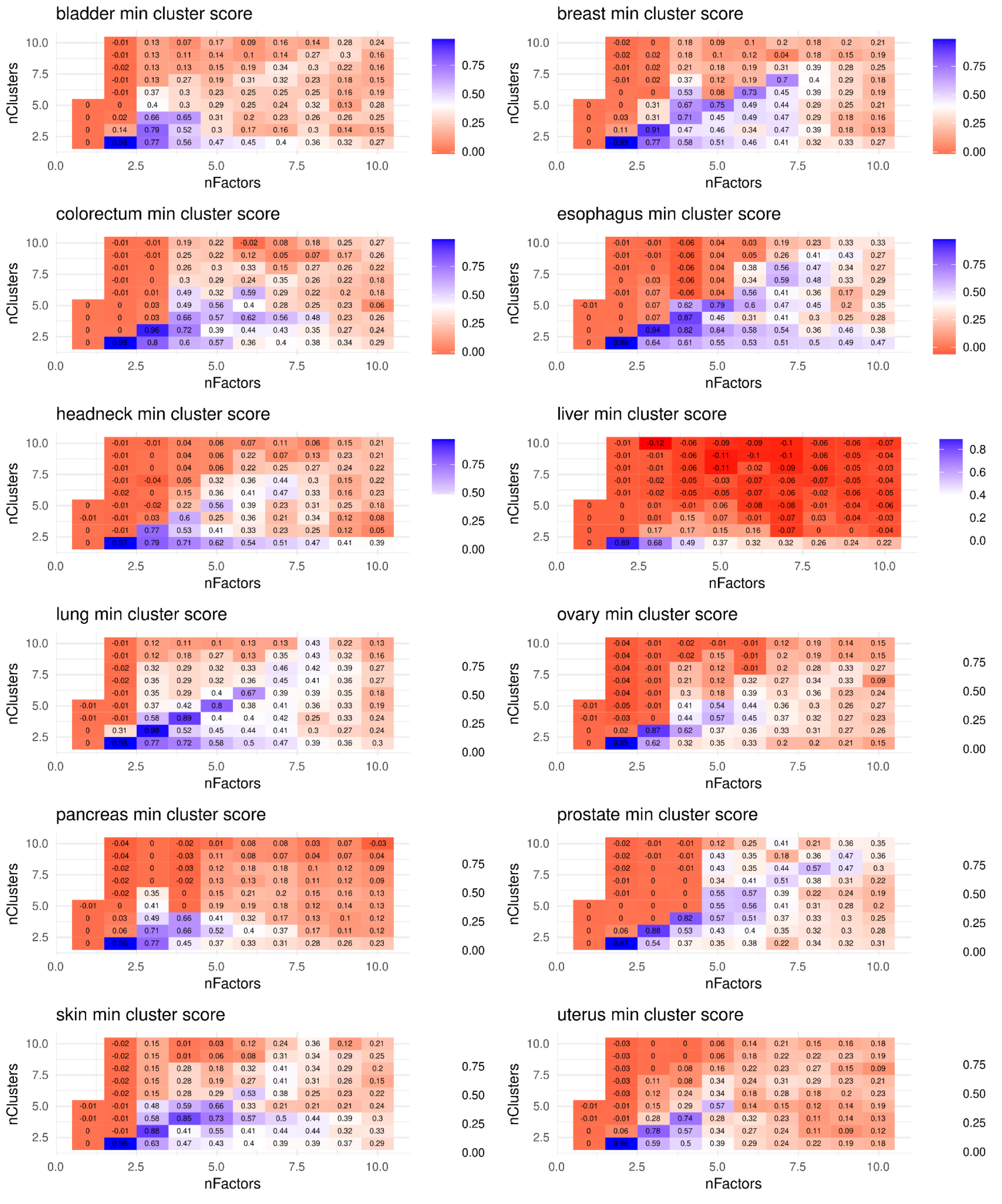


**Fig S8. Extracting RMD signatures within cancer types separately recovers multiple NMF factors.** Minimum silhouette index cluster score for different nFact and k values for the NMF methodology run in the 12 cancer types independently. Values selected are listed in Table S4.

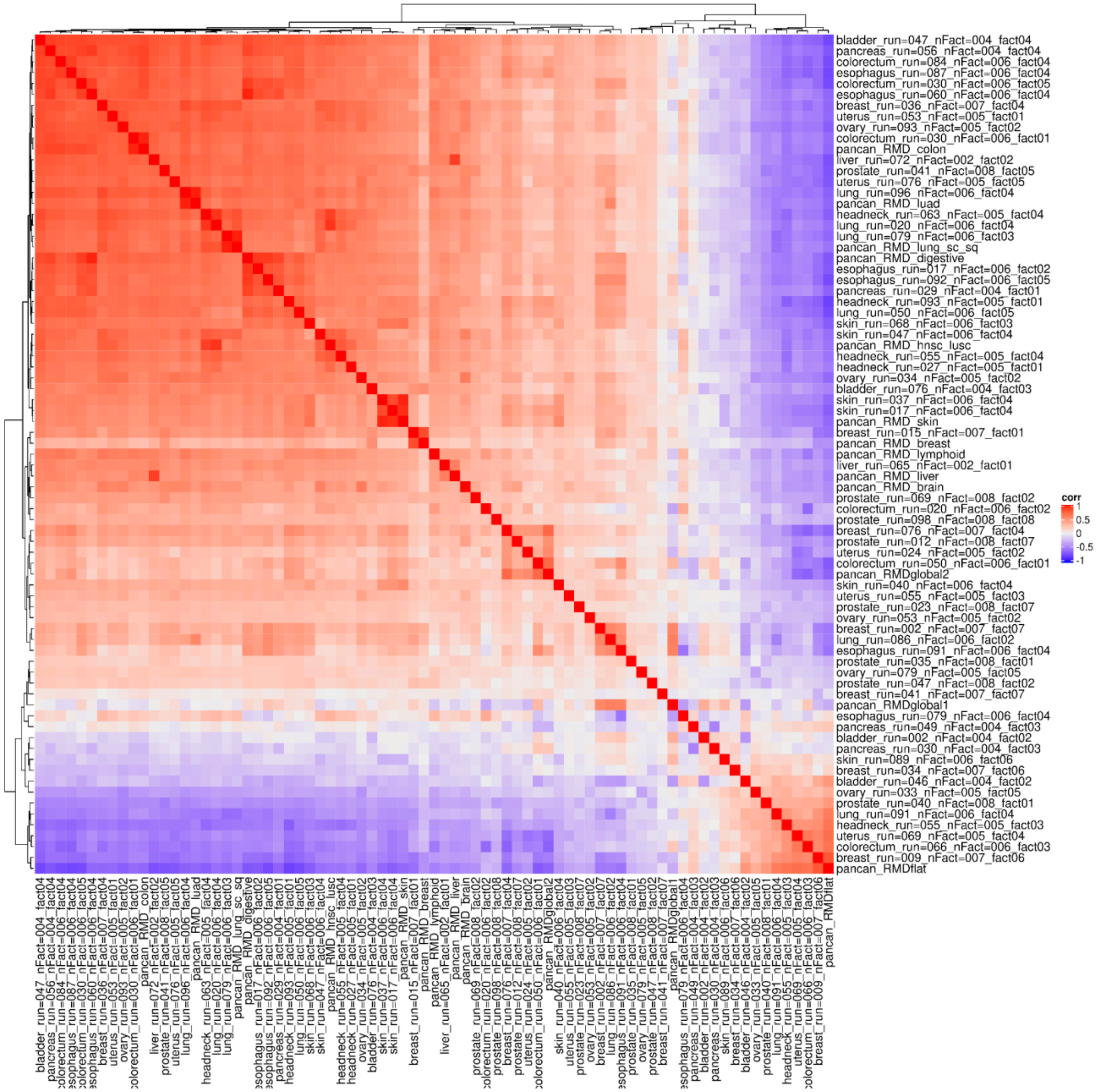


**Fig S9. Agreement between pan-cancer and individual cancer type RMD signatures.** Correlation between the RMDsignatures extracted in the pan-cancer NMF analysis (main results) and in NMF runs by cancer type (additional analysis) suggests that the 3 global RMD signatures (RMDflat, RMDglobal1 and RMDglobal2) can be recovered also from individual cancer types.

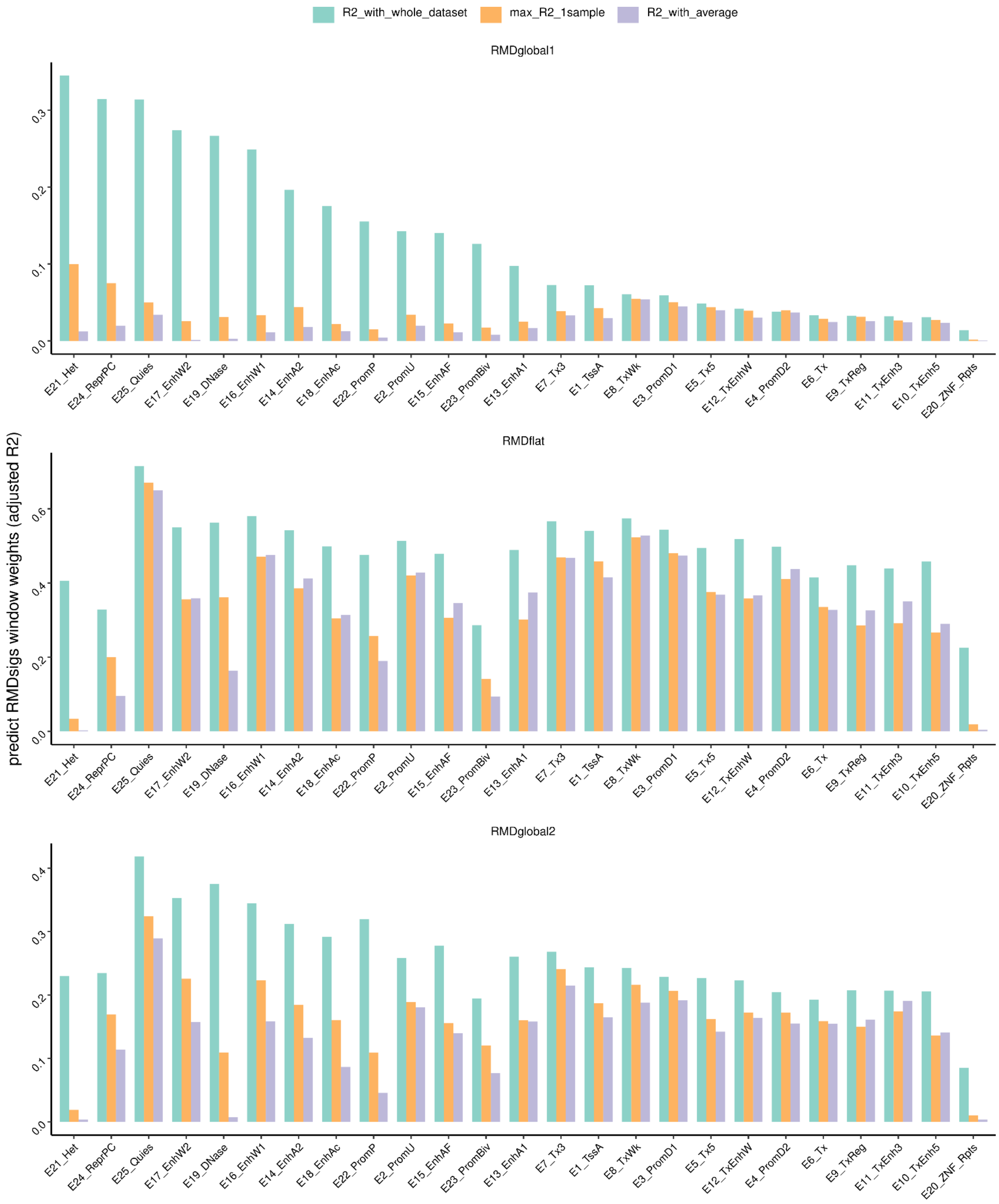


**Fig S10. Prediction of RMD signatures from the density of ChromHMM states.** Adjusted R2 of a regression predicting RMD signatures window weights from the chromHMM states density (% of the window cover by a particular state) (x axis) using the whole dataset, the average of the feature or selecting the maximum adjR2 of calculating it for each sample individually (max).

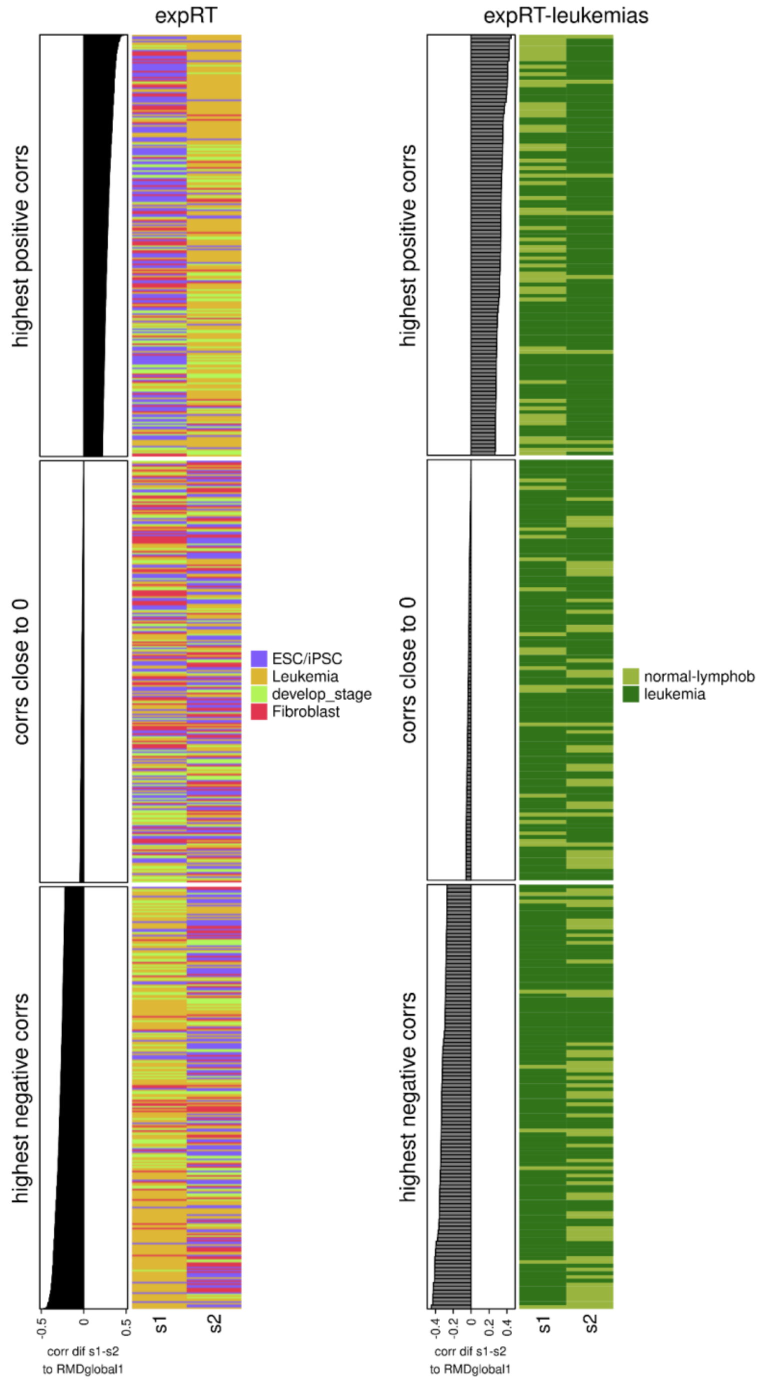


**Fig S11. Differences between RT profiles of samples correlates with RMDglobal1 window weights.** Correlation between RMDglobal1 signature and the difference between each pair of RT profiles. All combinations from the three data sets tested: experimentally measured RT (expRT) and a subset of expRT (lymphoblastoid cells and leukemias). Here we show the 1st (highest positive correlations), 5th (correlations close to 0) and 10th (highest negative correlations) deciles, ordered by correlation.

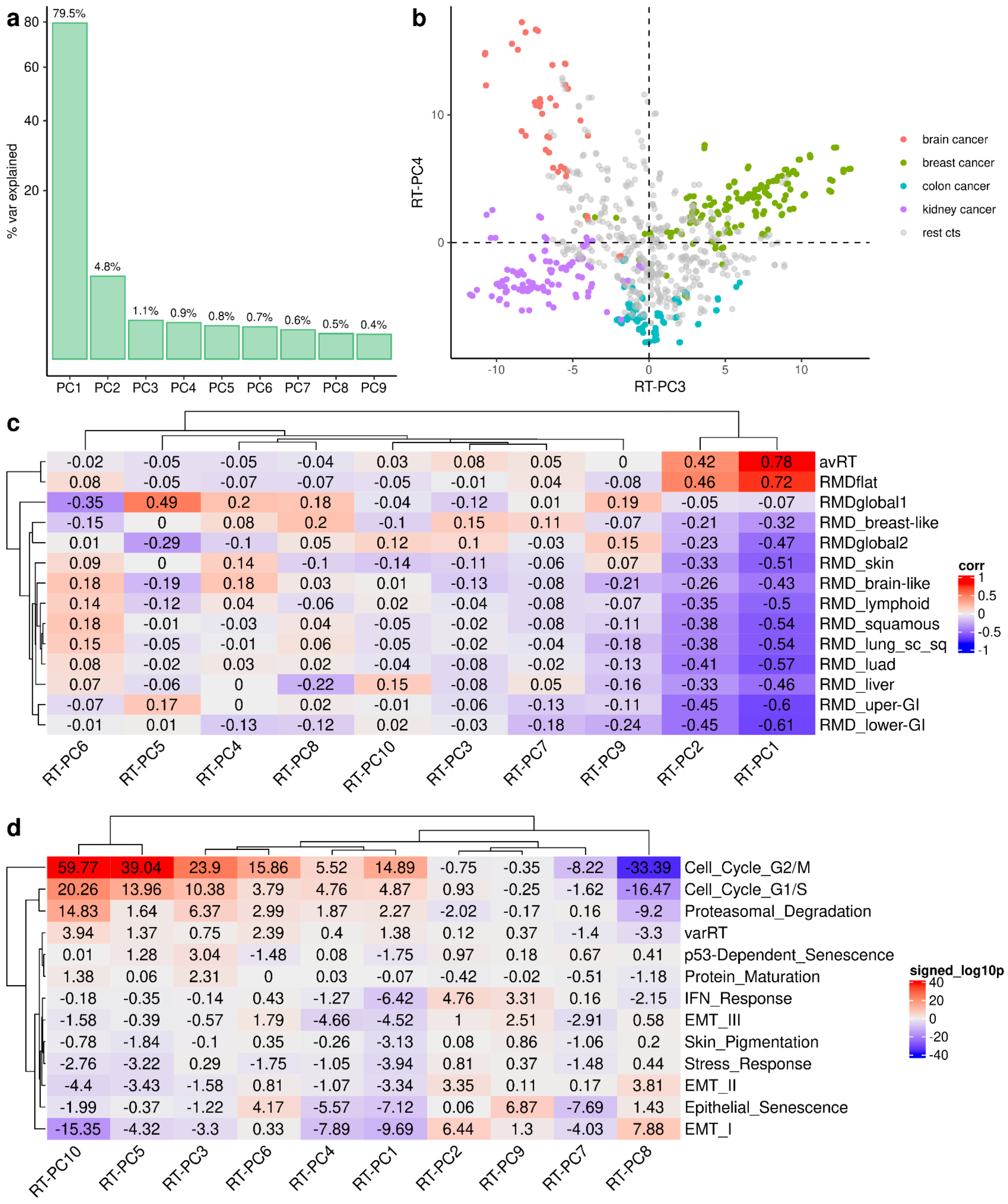


**Fig S12. A PCA in predicted RT from TCGA. a)** Variance explained by RT PC1 to PC9 from a PC analysis of predRT-TCGA dataset. **b)** RT-PC3 and RT-PC4 separate different cancer types: breast versus kidney cancer and brain versus colon cancer respectively. **c)** Pearson correlation between each RT-PC and the RMD signatures. RMDglobal1 highly correlates with RT-PC5. **d)** Associations (signed log10 p-value) between the GE in RHP pathways and the RT-PCs. We binarized RT-PC into high (3rd tertile) versus low (1st tertile).

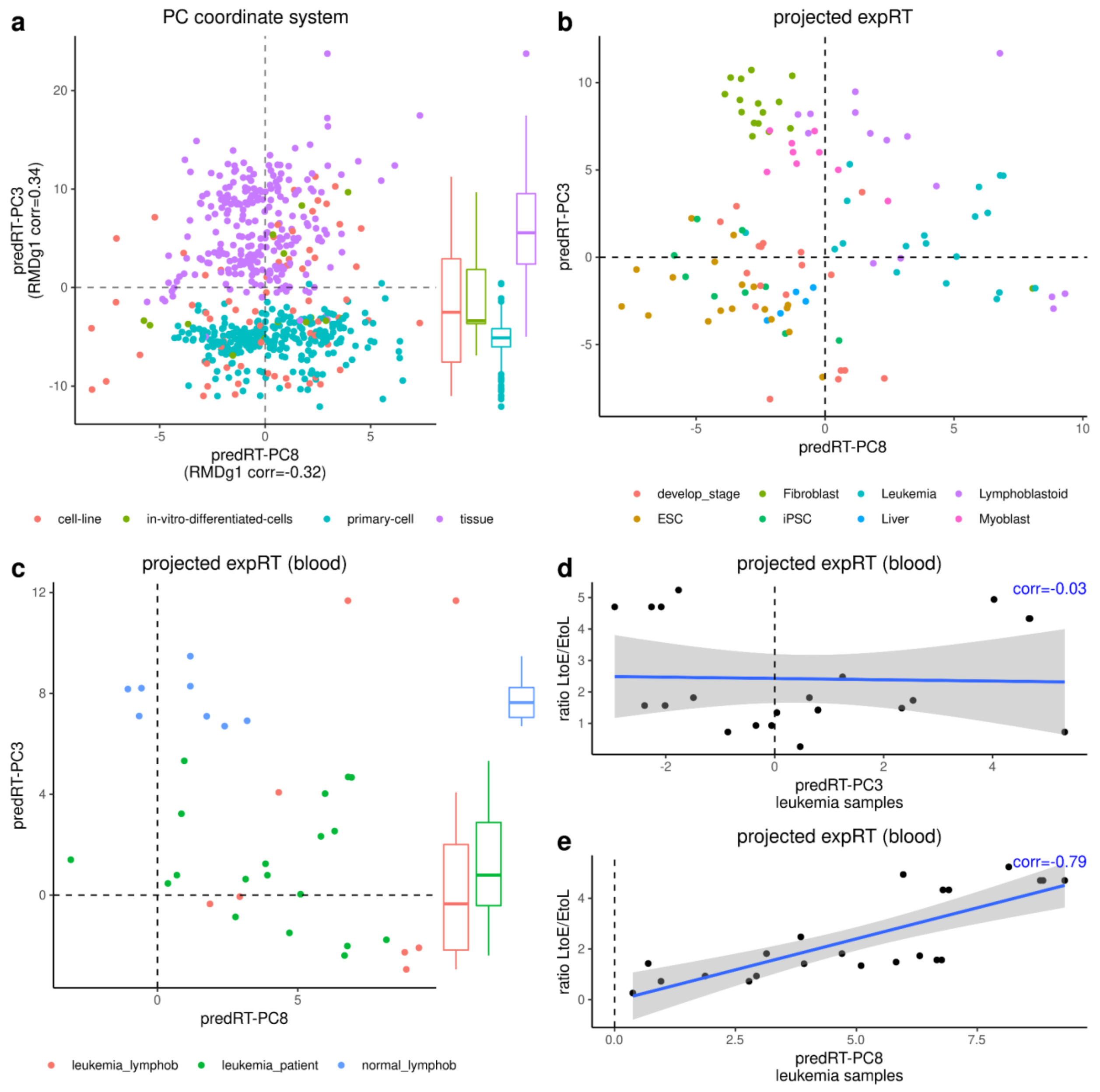


**Fig S13. A PCA to summarize the main trends in variation in the predicted RT (predRT) in the ENCODE dataset of various tissues, primary cells and cell lines. a)** PCs from predRT-ENCODE data with the highest correlation with RMDglobal1 signature, colored by cell type. PredRT-PC3 (RMDglobal1 R=-0.34) separates tissue versus primary cells. **b)** Projection of expRT data into the coordinate system of the PCA in predRT-ENCODE data. PredRT-PC3 separates healthy versus tumor in the projected expRT data. **c)** Projection of expRT data into the coordinate system of the PCA in a subset of predRT-ENCODE data encompassing lymphoblastoids cells and leukemia samples. **d-e)** Correlation between the projection of expRT leukemia samples in predRT-PC3 and 8 and the ratio of LtoE and EtoL changes from Ryba et al. 2012.

**a**

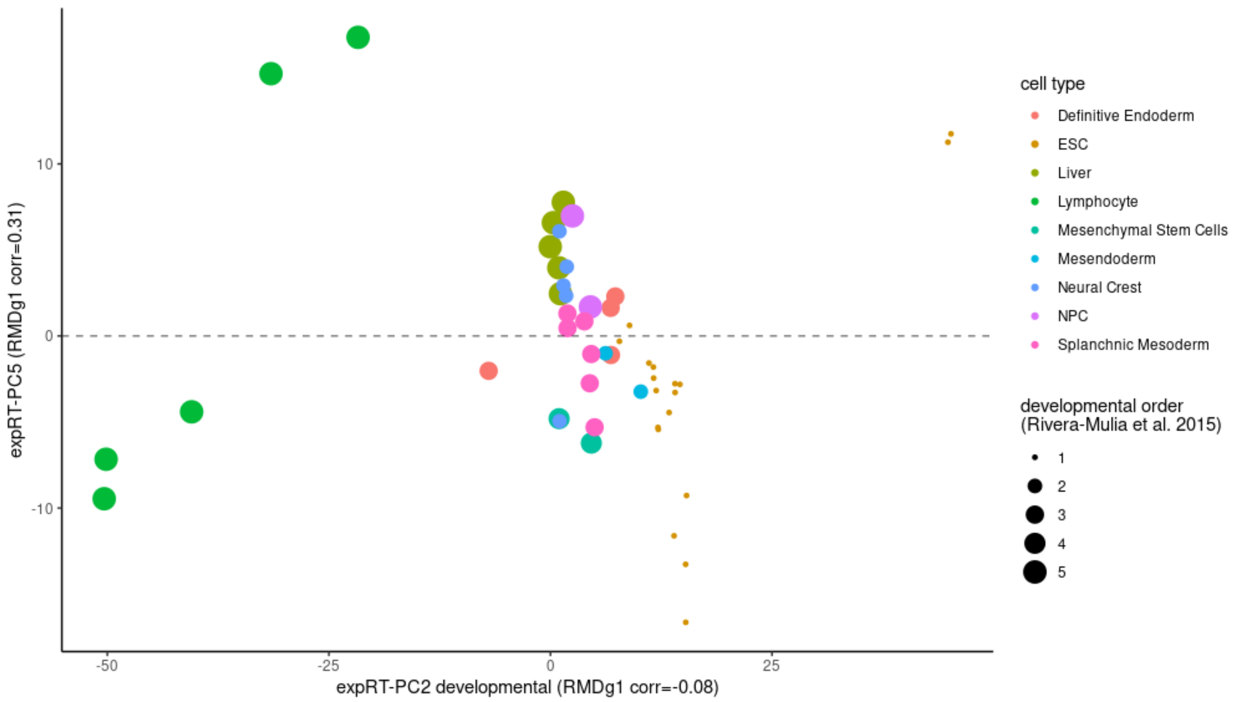


**b**

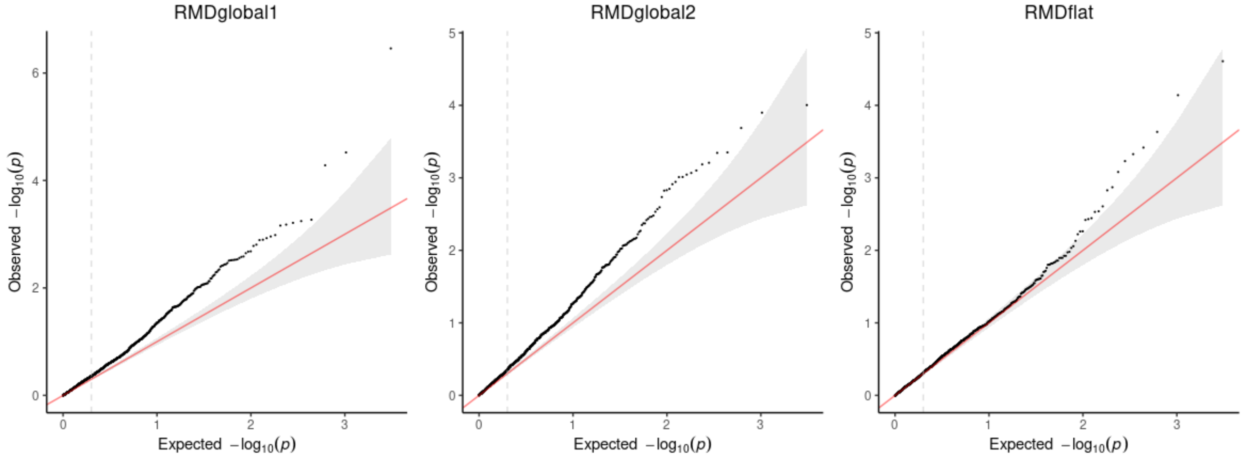


**Fig S14. a)** A PCA on the experimentally measured RT (expRT) profiles. ExpRT-PC5 correlates with RMDglobal1 mutation redistribution, while expRT-PC2 does not correlate with RMDglobal1, but is shown here since separates the developmental ordering (embryonal stem cells on the one end, versus developed lymphocytes on the other end, according to *Rivera-Mulia et al. 2015* classification). **b)** The quantile-quantile plots for p-values in association testing of three RMD global signatures with CNA in cancer genes and chromatin modifying genes.

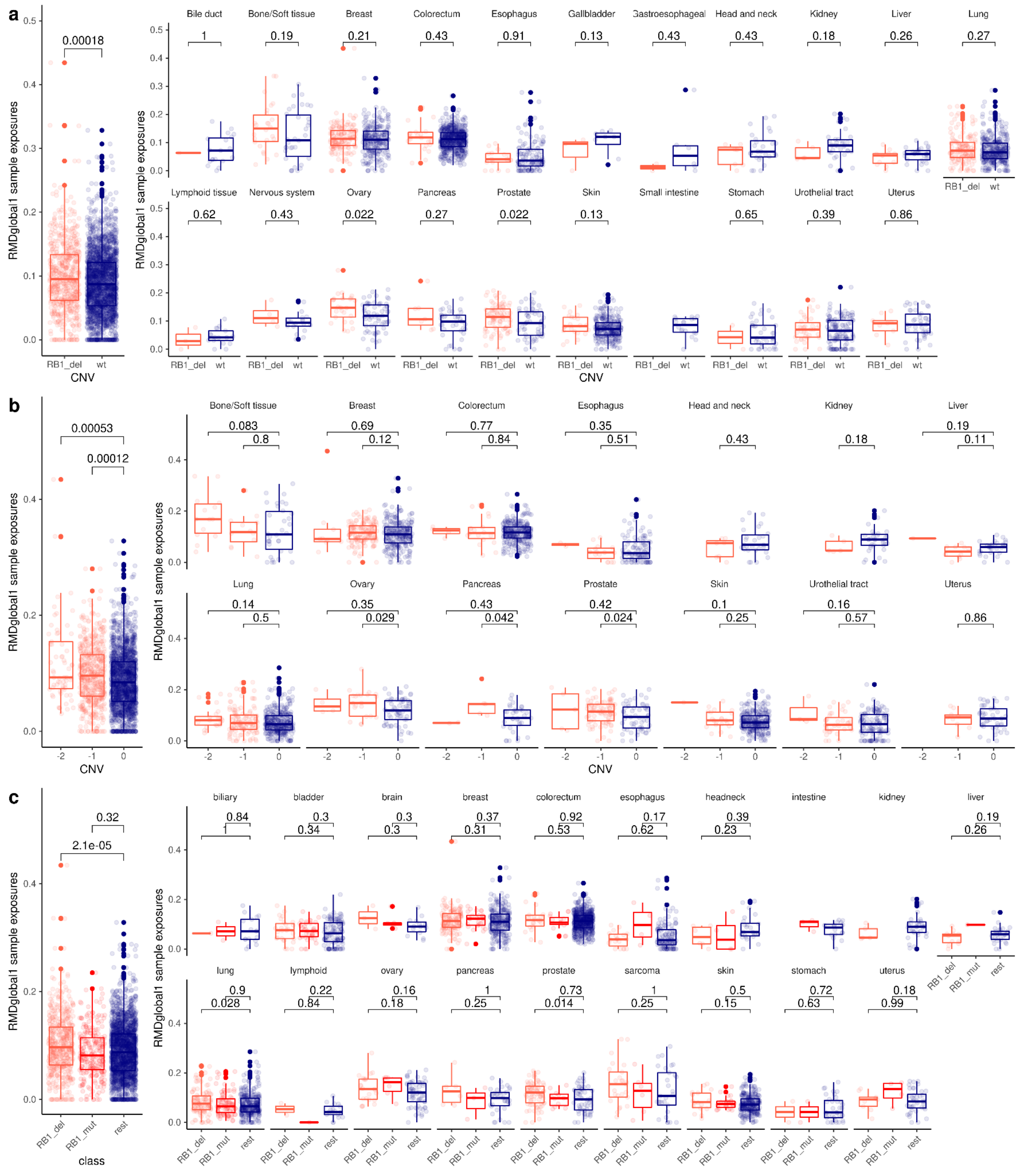


**Fig S15. Associations between RB1 deletion and mutation ("del" and "mut") and RMDglobal1 exposures. a)** Differences in RMDglobal1 exposures between RB1 deletion (-1 or -2 deletion) and wt pan-cancer (left) or by cancer type (right). **b)** Differences in RMDglobal1 exposures between 1 copy (-1) and 2 copies (-2) RB1 deletion and wt (0) pan-cancer (left) or by cancer type (right). **c)** Differences in RMDglobal1 exposures between RB1 deletion, RB1 mutations and wt pan-cancer (left) or by cancer type (right).

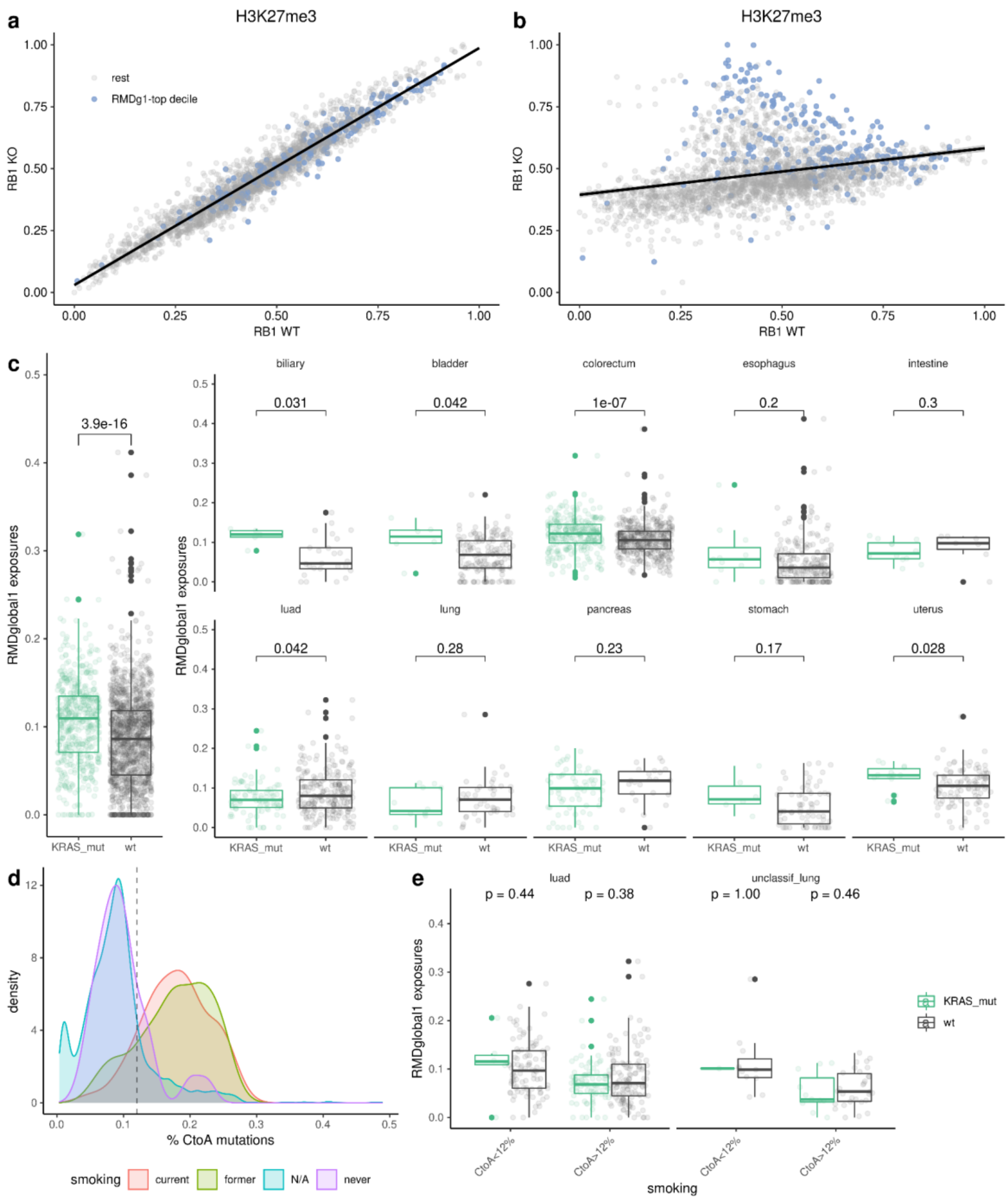


**Fig S16. Associations between *KRAS* mutation and RMDglobal1 exposures.** **a)** Correlation between the H3K27me3 heterochromatin profiles for samples with RB1 knock-out (“KO”) versus wild-type (“WT”). Each dot represents a window, colored by RMDglobal1 window weight top decile versus the rest of the windows. **b)** Correlation between the wild-type H3K27me3 and H3K9me3 profiles. Each dot represents a window, colored by RMDglobal1 window weight top decile versus the rest of the windows. **c)** Differences in RMDglobal1 exposures between *KRAS* mutated and wild-type in pan-cancer (left) or by cancer type (right); p-value by Mann-Whitney test. **d)** Distribution of fraction C>A mutations (proxy for tobacco smoking exposure) along the patients separated by their self-reported smoking status. We classified the lung cancer samples with missing data (“N/A” in the plot) according to the threshold C>A fraction of 0.12 (vertical line). **e)** Differences in RMDglobal1 exposures between *KRAS* mutated and *KRAS* wild-type lung cancers, after stratifying by putative smoking (C>A > 12%) and non-smoking (C>A < 12%) status, shows that *KRAS* mutation is not associated with RMDglobal1 in smokers, but might be in smokers (non-significant result, few non-smoker cancers are *KRAS* mutant).

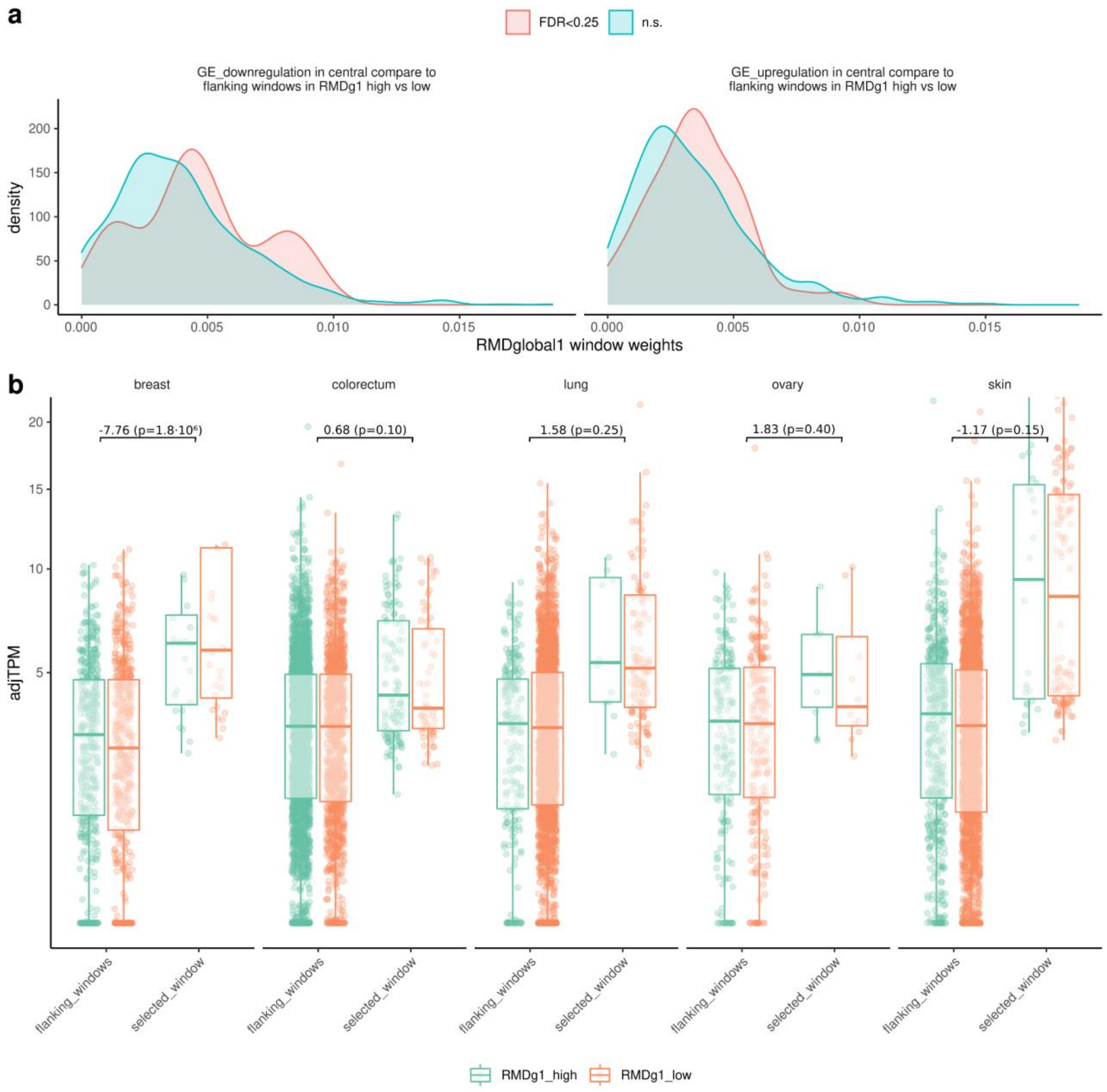


**Fig S17. Coordinated changes in gene expression associated with RMDglobal1 a)** Distribution of RMDglobal1 window weights for the windows detected as having coordinated gene expression changes (coordinated upregulation or downregulation) in one window, compared to their flanking windows, for RMDglobal1-high versus RMDglobal1-low tumors (p-value < 0.05) and the rest. **b)** Example of one selected window (chr2_144300002_145300001) and its flaking windows, split by RMDglobal1-high versus low tumor samples. This central window was selected because it showed significant gene expression downregulation.

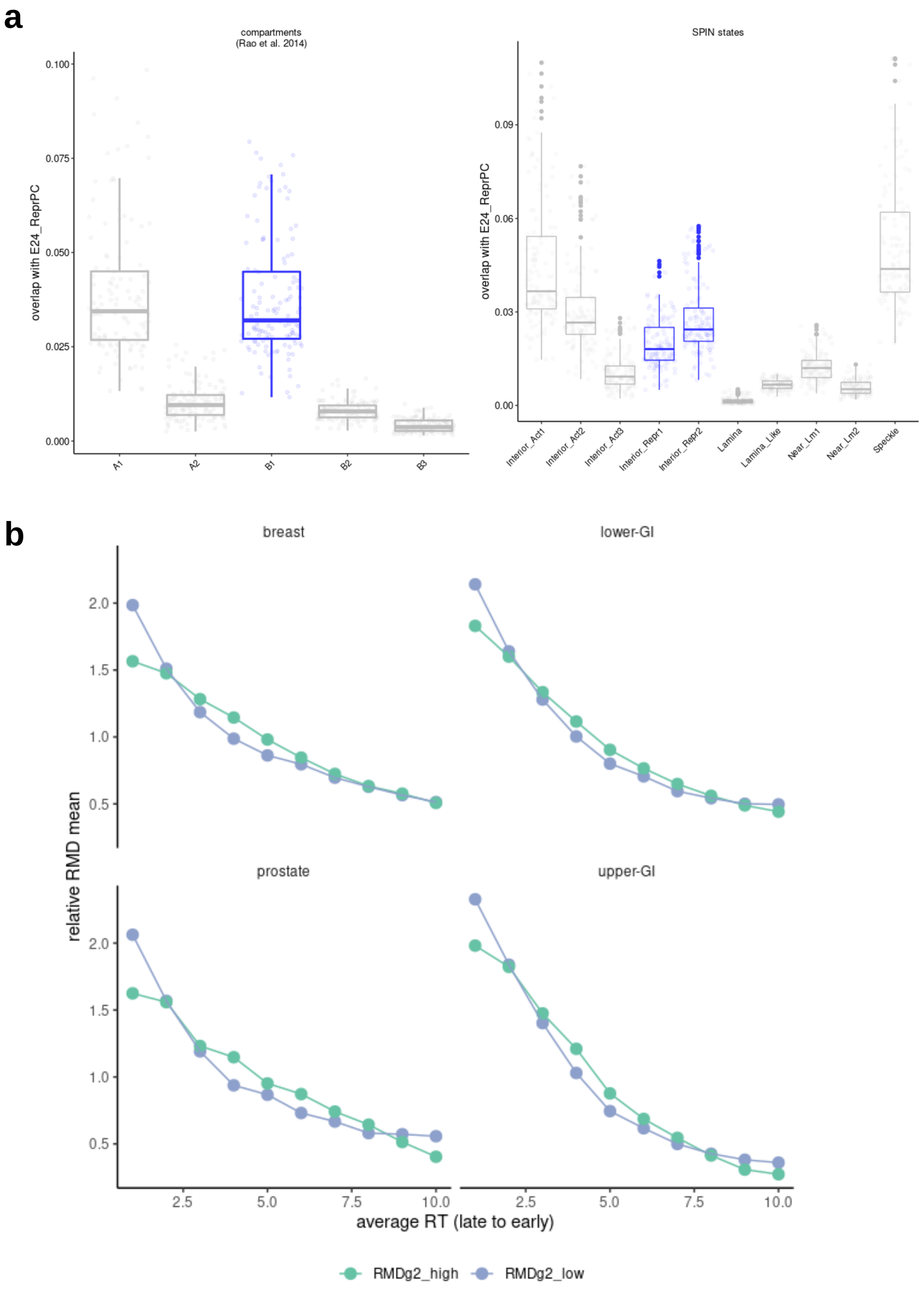


**Fig S18. a)** Association of various Hi-C based nuclear subcompartments with a polycomb-silenced chromatin state. The E24_ReprPC refers to the “ReprPC” (repressed Polycomb) state of ChromHMM analysis on Roadmap epigenome data, enriched with H3K27me3 histone mark. **b)** Regional mutation rates association with RT is altered by RMDglobal2 signature in various cancer types. Mean RMD values across 10 RT bins (late to early) for samples with RMDglobal2_high versus RMDglobal2_low across tissues.

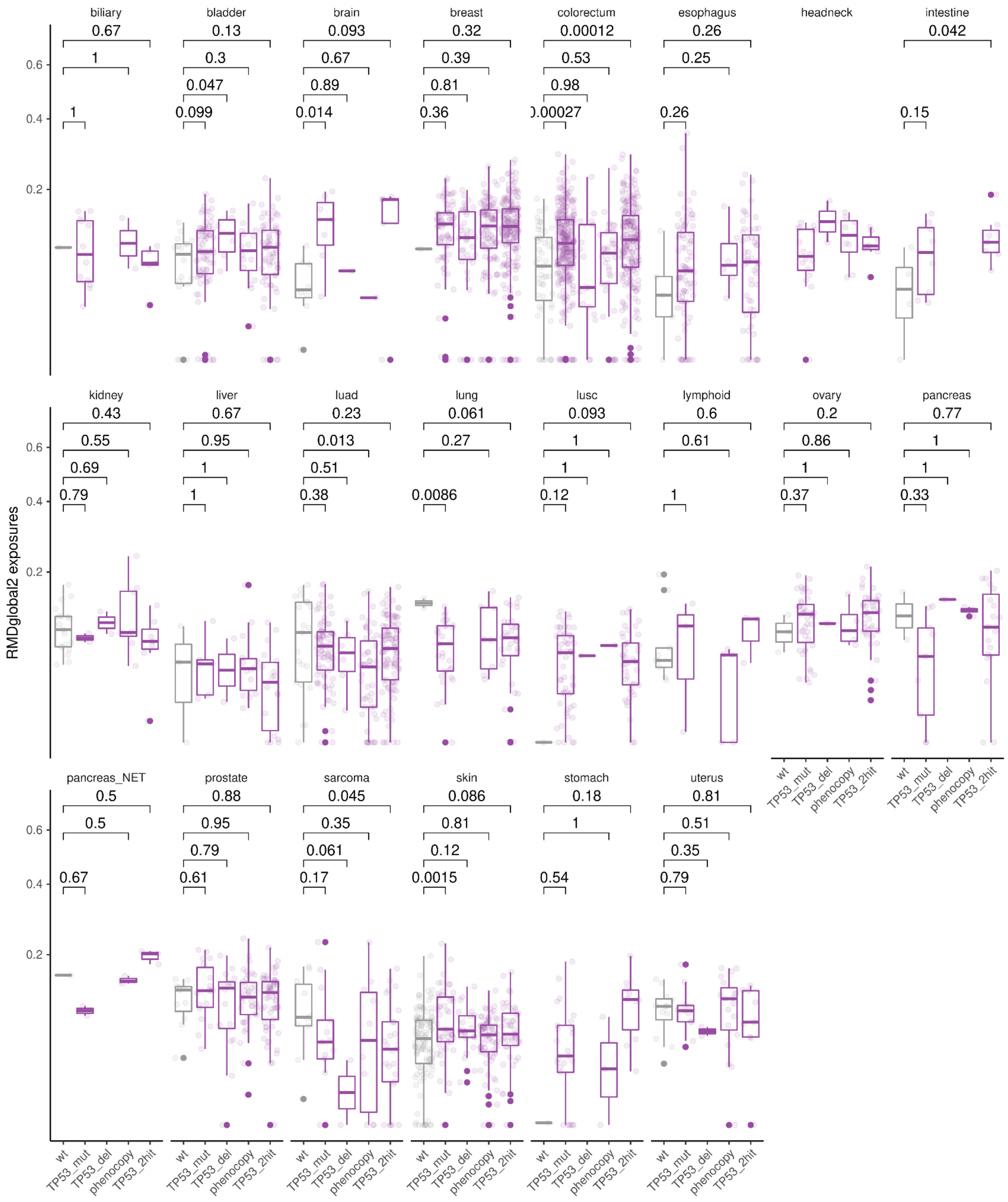


**Fig S19. Association of RMDglobal2 signature with various mechanisms of TP53 inactivation in different cancer types**. RMDglobal2 exposures grouped by: wild-type (wt), TP53 with 1 mutation (TP53_mut), TP53 with 1 deletion (TP53_del), TP53 phenocopied with an amplification in MDM2, MDM4 or PPM1D genes (TP53_pheno), or TP53 with two hits of the previously mentioned (TP53_2hit).

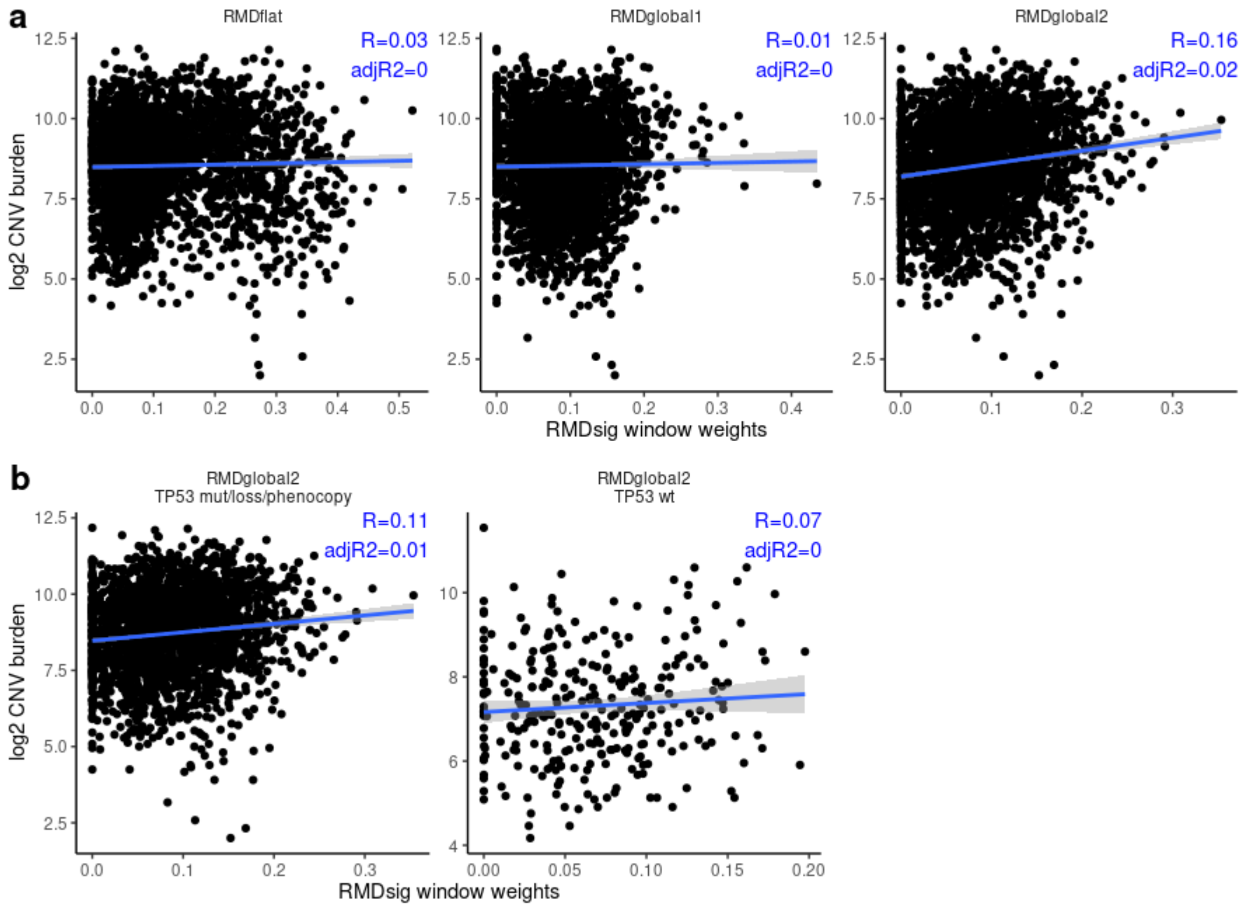


**Fig S20. Burden of copy number variant does not explain global RMD signatures. a)** Correlation between global RMD signatures and the CNV burden. **b)** Correlation between global RMDsignature2 and the CNV burden stratifying by TP53 functional status.
